# Steric-free bioorthogonal profiling of cellular acetylation and glycosylation via a fluorine-selenol displacement reaction (FSeDR)

**DOI:** 10.1101/2022.09.13.507737

**Authors:** Yue Zhao, Mi Zhao, Zhigang Lyu, Nicole Gorman, Todd R. Lewis, Aaron R. Goldman, Hsin-Yao Tang, Rongsheng E. Wang

## Abstract

Global detection and identification of protein post-translational modification (PTM) is a major bottleneck due to its dynamic property and rather low abundance. Tremendous efforts have been since made to develop antibody-based immunoaffinity enrichment or bioorthogonal chemistry-based chemical reporter approach but both suffer from inherent limitations. Following our previously reported steric-free tagging strategy, we hereby report the invention of selenol as a new generation of fluorine-displacement probe. The fluorine-selenol based displacement reaction enabled us to efficiently label and image acetylation and glycosylation at cellular level. We further pursued FSeDR in tandem with SILAC based quantitative proteomics to globally profile acetylation substrate proteins in a representative prostate cancer cell line PC3. Our results unraveled the fluorine-based toolbox for powerful chemical biology probing and allow for the future study of PTMs in a systemic manner.

## Introduction

Post-translational modifications (PTMs) add a variety of chemical functional groups to protein substrates after their biosynthesis.^1-2^ With more than 400 different types identified, most PTMs are dynamic, reversible, and regulated by a set of enzymes named ‘writers’, ‘erasers’, and ‘readers’, thereby mediating downstream signaling pathways and cellular events. Taking two representative common PTMs for instance: Lysine acetylation is a small but essential PTM that is associated with metabolism, signal transduction, gene transcription, cell differentiation, and apoptosis, etc.^3^ The acetylation on protein is written by lysine acetyltransferases (KATs) which can transfer an acetyl group from acetyl-coenzyme A (Ac-CoA) to the ε-amino group of lysine residue and are grouped into three families: Gcn5-related N-acetyltransferases (GNATs, including GCN5, PCAF, HAT1), MYST (<u>M</u>OZ, <u>Y</u>bf2, <u>S</u>as2, and <u>T</u>ip60), and p300/CBP.^4^ Glycosylation is a abundant and diverse PTM that promotes the folding, stability, trafficking of proteins.^5-6^ Cell surface glycoproteins help mediate cell adhesion, ligand binding, cell-cell communication, and immune responses.^7^ The enzyme-mediated glycosylation is driven by glycosyltransferases^8^ that utilize activated nucleotide sugars as glycosyl donors and finally attach glycans to proteins mainly in two ways: N-linked glycosylation to asparagine (Asn) and O-linked glycosylation to serine (Ser), threonine (Thr) or tyrosine (Tyr). N-linked glycans all contain a core pentasaccharide of three mannose and two N-acetylglucosamine (GlcNAc),^9^ while mucin-type O-glycosylation is attached to Ser or Thr residues through a N-acetylgalactosamine (GalNAc). Dysregulation of PTMs have been shown pivotal towards the onset and relapse of human diseases. Increased sialylation on cancer cell surface antigens promotes cell detachment from the tumor mass.^10^ Other changes in glycosylation patterns are linked to cancer,^11^ chronic kidney disease, and neurodegenerative diseases.^12^ Similar to glycosylation, emerging evidences indicate non-histone proteins as the major portion of the acetylome.^13 14-15^ Dysfunction of acetylation is associated with many diseases such as type 2 diabetes,^16^ Tau-mediated Alzheimer’s progression,^17^ cancer migration^18-19^ and so on.^20^

Detecting these PTMs and understanding the substrate proteins would be of urgent importance for diagnostic and therapeutic purposes but were largely limited by the currently available tools.^21^ Classical methods to detect and analyze PTMs rely on antibodies and antibody-based LC-MS/MS proteomics analysis.^22^ However, the small-sized PTMs makes their recognition and enrichment challenging and the results varied significantly per antibody.^21, 23^ With a cocktail of multiple monoclonal antibodies directed towards acetyl lysine, only 18% shared sites were observed.^23^ Similarly, the use of lectins and antibodies for glycoprotein enrichment is limited by the low substrate binding affinities and/or poor specificities, particularly for O-glycans.^24-25 26^ Alternatively, chemical biology approaches with the bioorthogonal chemical reporter incorporated PTM cofactor or metabolic precursor have been widely adopted. The classic azide or alkyne tag can be derivatized into acetyl-CoA to label and identify acetylation substrates,^27-28^ and into monosaccharides for label and identify glycosylated proteins as metabolic glycan engineering.^29^ Followed with copper-catalyzed azide-alkyne cycloaddition (CuAAC) reaction, one can expend these tags with functional groups such as fluorophores for detection or biotin for pull-down and chemical proteomics studies (Figure 1). Yet, the alkyne or azide tags were recently revealed to be bulky in size,^21^ and thereby cannot be incorporated by most acetyltransferases^30^ or even accommodated well by glycosyltransferases.^31^ While mutations in protein pockets can be pursued, ^30, 32-33^ the tedious process may compromise the correct folding of PTM ‘writers’ or even further perturb the intrinsic acylome complex.

**Figure 1.**
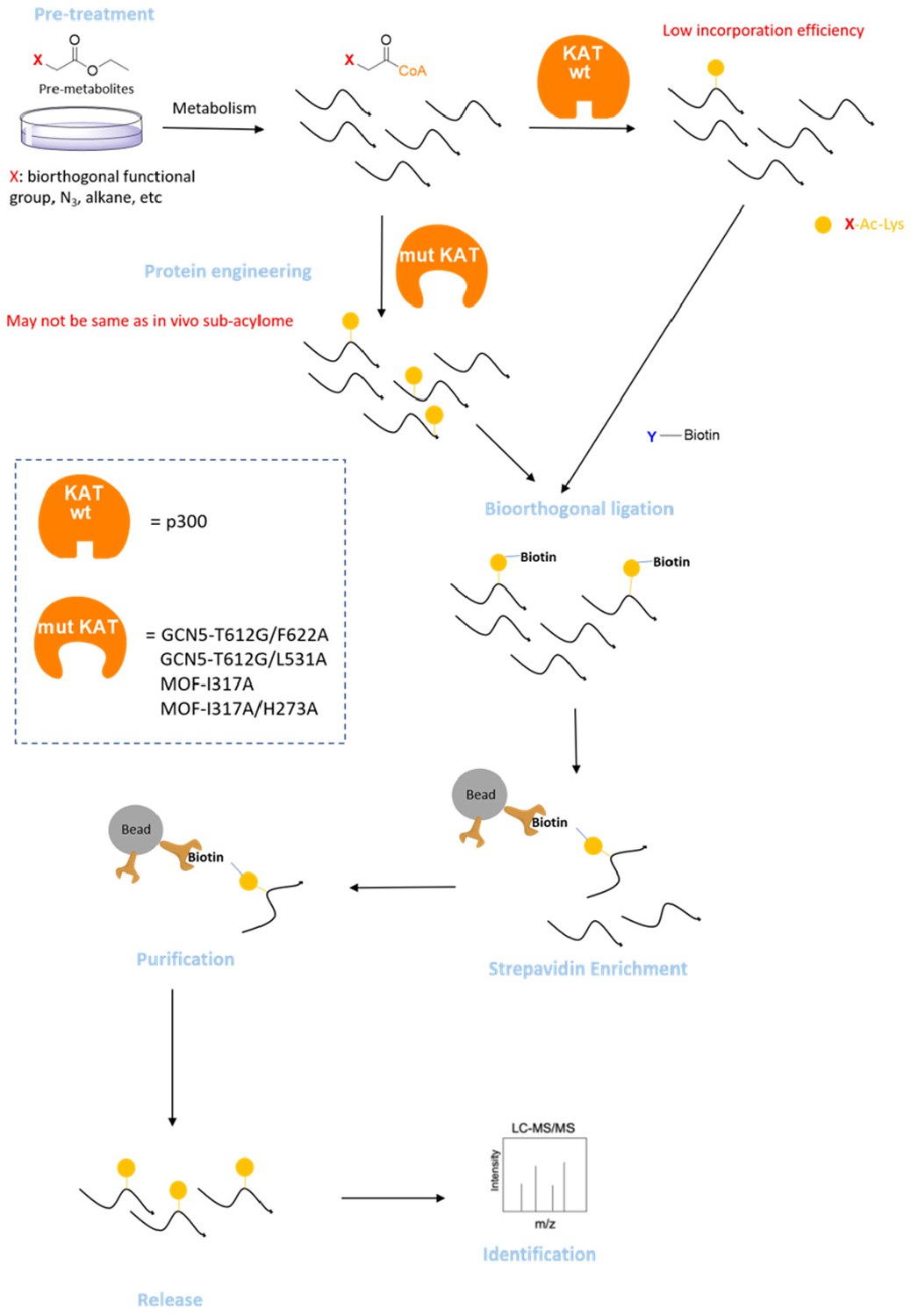
General procedures for chemical proteomics study of PTM substrate proteins. Acetylation is used here as an example. KAT: lysine acetyl transferase (p300, GCN5, MOF); wt: wild type; mut: mutant.

Previously, our lab invented a steric-free fluorine tagging strategy and demonstrated the use of aryl thiol probes to initiate a bioorthogonal fluorine-thiol displacement reaction (FTDR), which enables superior universal labeling of acetylation proteins in mammalian cell lines to the traditional CuAAC-based alkyne/azide labeling.^21^ Nevertheless, the prototype reaction requires a mildly basic pH environment (8.5) and has a slow reaction kinetics. Herein, we report the creation of selenium-based selenol probes which leads to fluorine-selenol displacement reaction (FSeDR) that has improved rate constant and occurs at neural pH 7.2. With the selenol probes, we efficiently labeled the cellular substrate proteins of acetylation and glycosylation, and for the first time demonstrated the tandem use of quantitative proteomics to identify 43 new acetylation substrate proteins. Some validated proteins such as DDX54 and HSPA6 may reveal significant regulation roles that acetylation plays in the related cellular signaling.

## Results and Discussion

### Synthesis and characterization of the selenol probe

Compared to the previously exploited sulfur atom (Figure 2A),^21^ most selenium-hydrogen bond is weaker and the selenols’ pKa was lower (Figure 2B). The previously-used aryl thiol probe (2,4,6-trimethoxybenzenethiol) has a pKa of 6.7, suggesting that the preferred mildly basic pH of the thiol probes is likely due to the need for more nucleophilic thiolate formation. The lower pKa of selenols ensures that they can be fully deprotonated at neutral pH to become selenolates, which have higher polarizability and are one order of magnitude more nucleophilic than thiolates (Figure 2C).^34^ We therefore synthesized the two model selenol compounds (Alkyl-SeH, **13** and Ar-SeH,**14**) according to Scheme S1 (Supporting Information). The previously reported thiol compound **12** was prepared as a control.^21^ Reaction of each compound with the model F-acetamide substrate **11** was carried out in Tris buffer for 4 h, followed by LC-MS analysis to compare the conversion yield according to the integration of peak areas (Figure 2D). In general, selenol compounds displayed higher efficacy than the thiophenol compound. The similar yield caused by Alkyl-SeH and Aryl-SeH are likely due to the faster dimerization of the aromatic selenol, which was observed to occur quickly by LC-MS. Thus, we decided to focus on alkyl selenols in this study.

**Figure 2.**
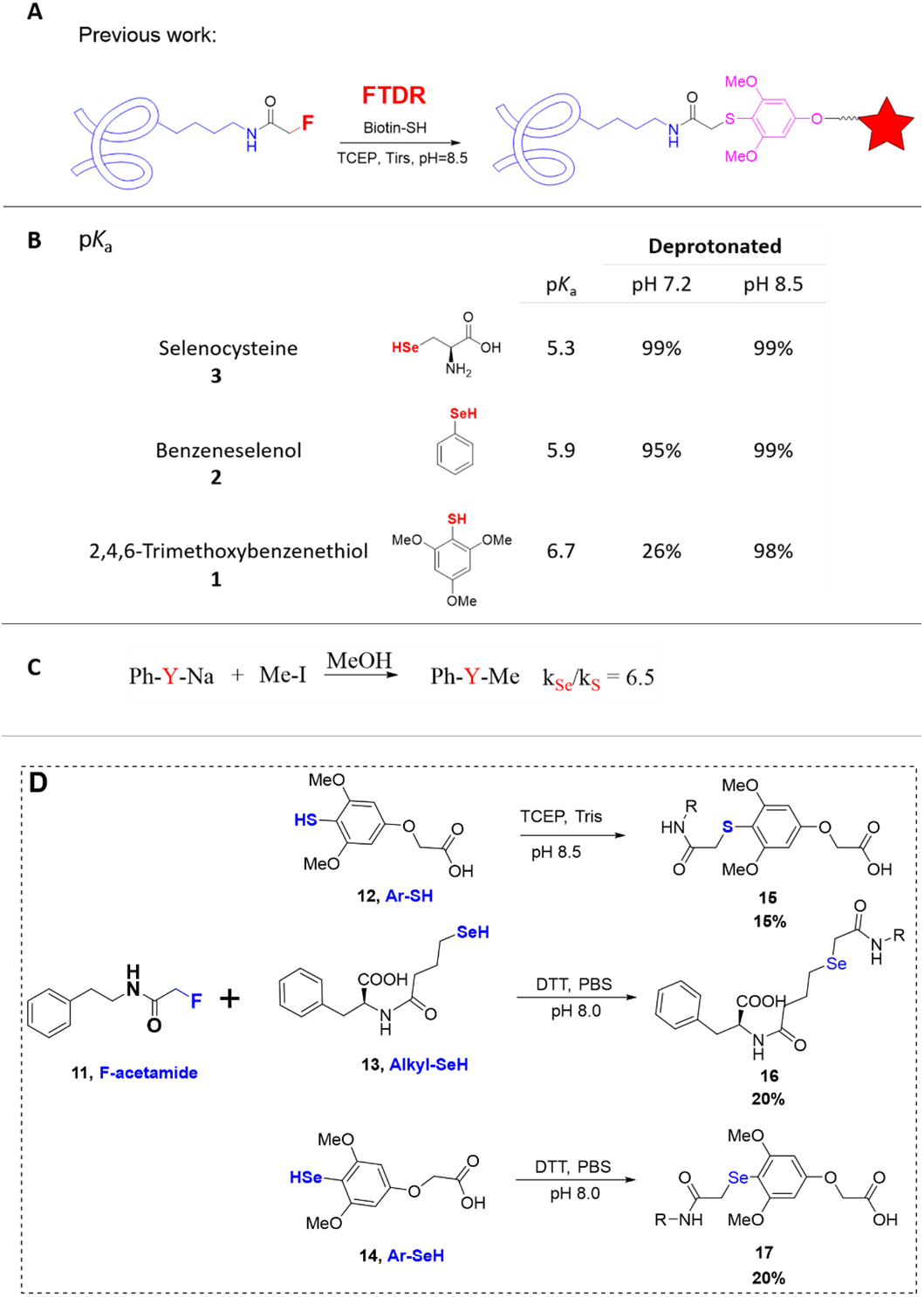
Model selenol probes for fluorine-displacement reactions. (A) Our previously reported fluorine-thiol displacement reaction (FTDR) (B) pKa value for 2,4,6-trimethoxybenzenethiol (**1**), benzeneselenol (**2**), and selenocysteine (**3**). The data was referred from the SciFinder database. Based on the Henderson-Hasselbach equation: pH = pKa + log [A-]/[HA], the deprotonated ratio was calculated as [A^-^]/([A^-^]+[HA]). (C) Comparison of selenium and sulfur nucleophilicity in a model S_N_2 substitution reaction using phenyl selenolate/thiolate with methyl iodide. (D) Comparison of benzenethiol and selenol probes regarding their displacement of fluorine from the model substrate F-acetamide. The reactions were performed in parallel for 4 h.

To protect the highly reactive selenol warhead from dimerization or other oxidations, we figured out to store the selenols with the cyano protecting group as the stable intermediate **10** (Scheme S1) and would only deprotect it right before use. The best deprotection condition was screened by testing model probe **10** with sodium cyanoborohydride (NaCNBH_3_) and sodium borohydride (NaBH_4_). It turned out NaCNBH_3_ cannot deprotect cyano group even with higher concentration and longer incubation times. On the contrary, 5 equivalent of NaBH_4_ can remove cyano group very efficiently in just 10 min (Figure S1A-S1C, Supporting Information). The selenol probe is also readily to form dimer under exposure to air which will block the reactive selenium anion warhead.^35^ To reduce the dimer, we explored tris(2-carboxyethyl) phosphine (TCEP), dithiothreitol (DTT), NaCNBH_3_, or NaBH_4_ to break diselenide bonds (Figure S1D-S1H). Although being odorless, more powerful and resistant to oxidation in air, TCEP was unfortunately found to react with selenol in a short time (less than 30 min). The radical deselenization reaction formed the unwanted TCEP=Se product.^36^ In the end, we found that 25 equivalent DTT is the optimal condition to reduce the selenol dimer and to maintain the probe in its monomer form for several hours at low concentration (less than 10 mM).

Next, we titrated the optimal pH range for this fluorine-displacement reaction (Figure 3A-3D), and observed that the reaction rate significantly increased with pH values until the pH reached 7.2. Under the physiological condition, pH 7.2, the reaction can reach 86% reaction conversion within 12 h. The product **16** was purified out and the structure was confirmed by NMR study. At higher pH values such as 8.0 (Figure 3D), transformation of reactive monomer probe into dimers became dominant and slowed down the reaction conversion. We attempted the reaction kinetics studies (Figure 3E) at concentrations of 5 mM and 10 mM, respectively, and the resulting 2^nd^ order rate constant is ∼ 4.25 ± 0.18 × 10^−3^ M^-1^s^-1^, which is at least 4-5 times faster than the previously reported FTDR reaction.^21^

**Figure 3.**
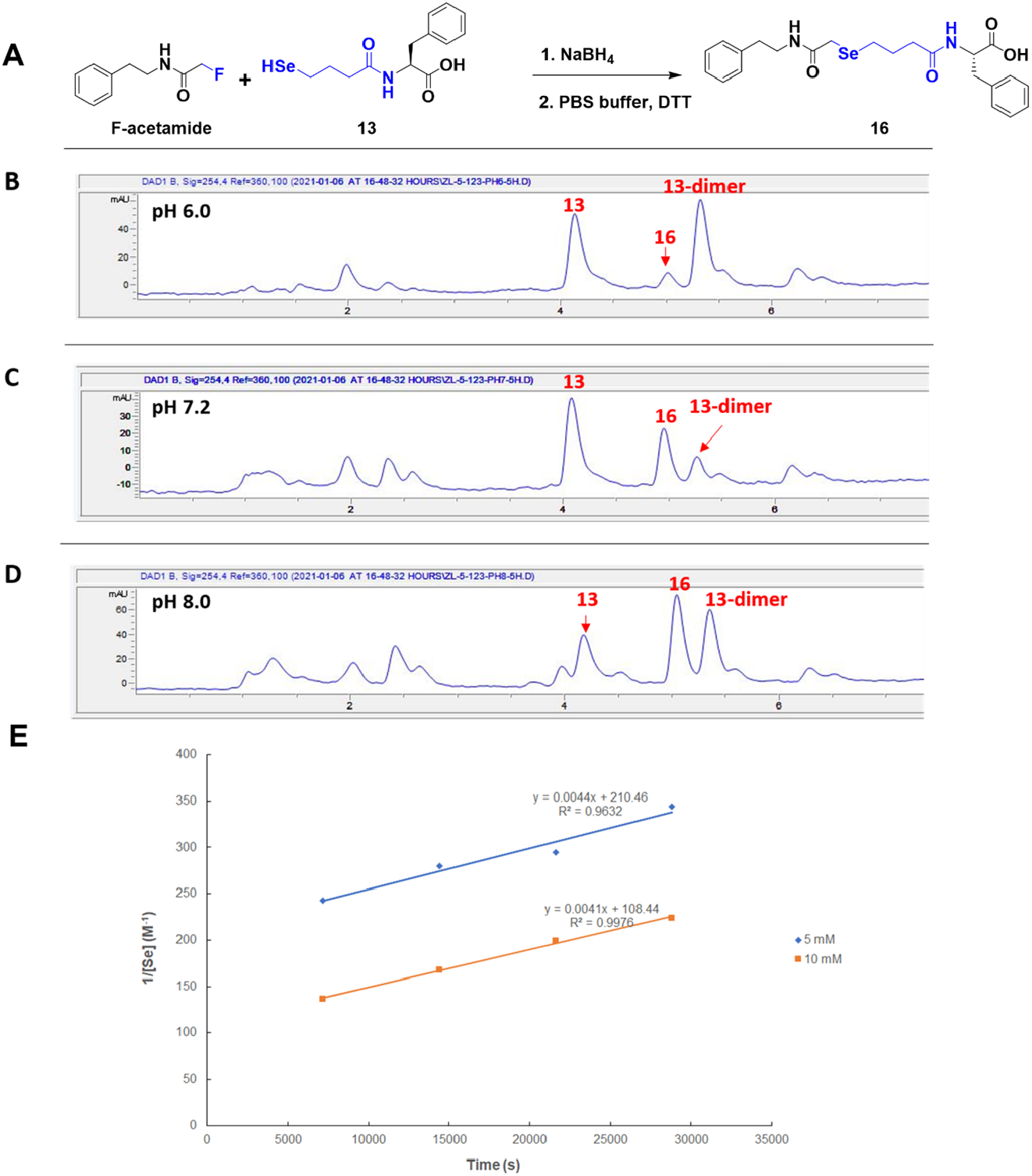
Characterization of the fluorine-selenol displacement reaction (FSeDR). (A) Reaction scheme for model FSeDR, with the use of model nucleophile probe **13** and F-acetamide substrate **11**. (B-D) pH titration of the FSeDR: representative LC-MS spectra of the model reaction after 5 h of incubation under varied pH’s. (E) The 2^nd^ order reaction kinetics measurement of the model FSeDR at pH 7.0. The average rate constant is (4.25 ± 0.18) × 10^−3^ M^-1^ S^-1^.

### Labeling and pull down of the model protein using FSeDR

With the new selenol probes, we set out to test their labeling of fluorinated model protein, bovine serum albumin (BSA). The desthiobiotin probe was constructed (**19**, Biotin-SeH, Scheme S2, Supporting Information) which contains selenol as the warhead and the PEG2 linker as the connecting unit to improve probe solubility. The final probe is very soluble in water and can be used to prepare scaled probe stock solution with concentrations up to 100 mM. Next, we examined the FSeDR labeling on the fluoroacetylated model protein BSA by incubating the Biotin-SeH probe with protein for varied time periods, followed with subsequent in-gel fluorescent imaging to detect the presence of biotin tag on proteins (Figure 4A-4B). As a control, another set of fluoroacetylated BSA was incubated with the published Biotin-SH probe^21^ and analyzed in parallel. As a result, the treatment groups using Biotin-SeH has generated efficient biotinylation on BSA, as evidenced by significant and saturating fluorescent signals within 3h. On the contrary, treatment with Biotin-SH probe rendered much weaker signals with biotinylation gradually reaching saturation after 6h of reaction. The minimal signal observed from the control group (unmodified BSA) indicated the relative specificity of Biotin−SeH/SH (Figure 4B).

**Figure 4.**
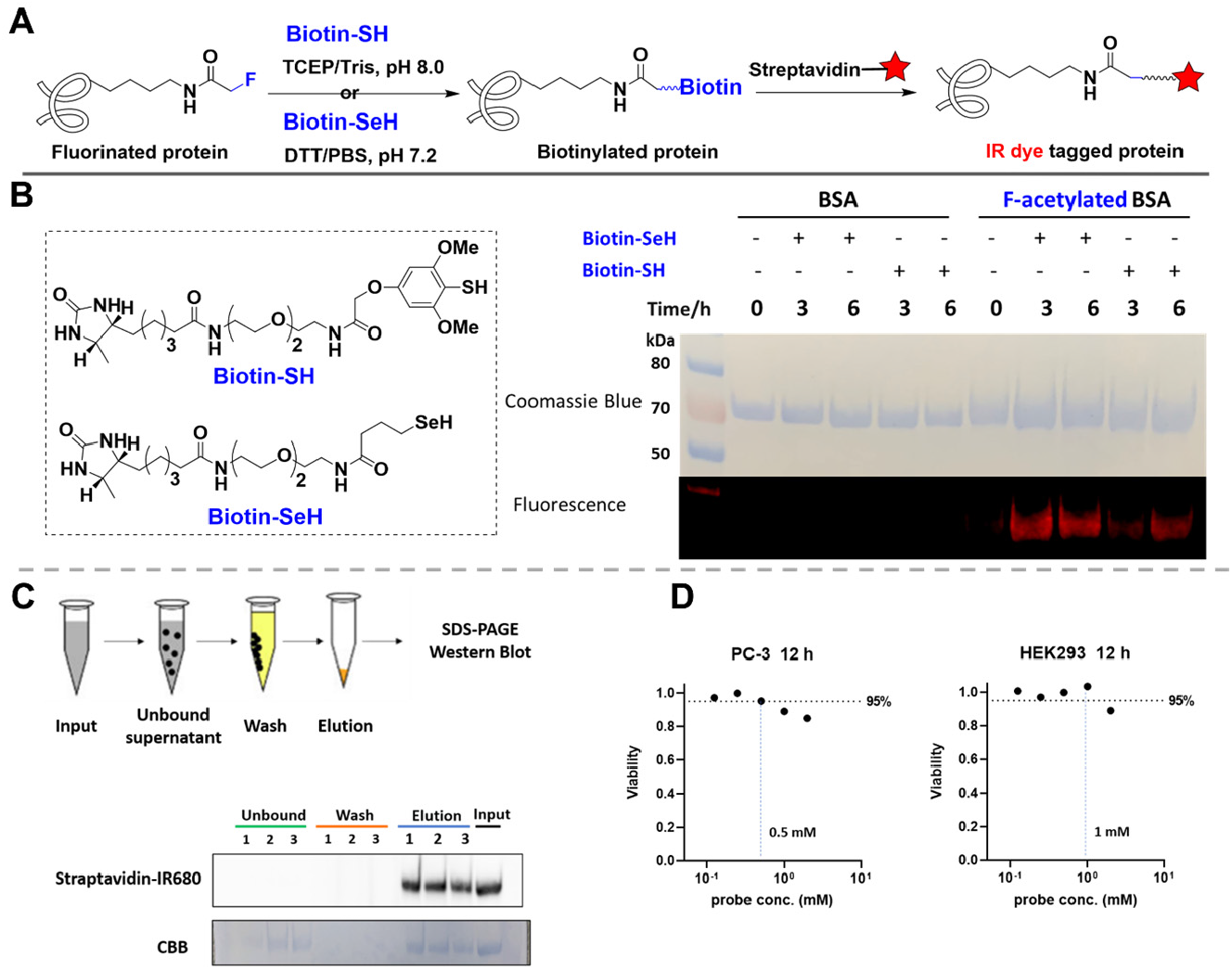
Labeling and pull-down of the model BSA protein using Biotin-SeH probe. (A) The experimental scheme for protein labeling with the adoption of different biotin probes and the respective reaction conditions. (B) *In vitro* labeling of BSA protein using Biotin-SH (pH 8.5, Tris buffer) or Biotin-SeH probes (pH 7.2, PBS buffer). The fluorescent image is for in-gel detection of IR dye-conjugated streptavidin that is incubated with BSA after labeling reaction. F-acetylated BSA was prepared by random lysine conjugation of BSA with F-acetyl NHS ester. (C) Experimental scheme for enrichment of biotinylated BSA generated by FSeDR labeling. With the streptavidin magnetic beads, the effect of detergent was tested. Condition 1: DPBS only; condition 2: DPBS containing 0.1% SDS; condition 3: DPBS containing 0.2 % SDS. (D) Cytotoxicity evaluations of the Biotin-SeH probe in live cancer cell lines (PC-3, left; HEK293, right).

Moving forward, we attempted the pull down of BSA model protein from buffer using our probes. As shown in Figure 4C, the biotinylated BSA proteins based on FSeDR were enriched thoroughly by streptavidin-magnetic beads. Inclusion of 0.1-0.2% SDS in the solution could efficiently reduce non-specific protein binding while still retaining the desired biotin-labeled target proteins. To further gauge the Biotin-SeH probe’s potential, we evaluate its cytotoxicity in mammalian cell lines (Figure 4D) and found that PC-3 and HEK293 both exhibited good tolerance of the selenol probe, with more than 85% viability after 12h incubation.

### Labeling and pull down of cellular proteins using FSeDR

Next, we applied the Biotin-SeH probe towards live HEK293 cells, in order to evaluate whether FSeDR labeling could be practically utilized to globally enrich protein substrates. Treatment with ethyl fluoroacetate at the first step would allow the cells to convert the ethyl fluoroacetate to F-Acetyl CoA.^21^ The cells were then lysed and incubated with the probe in step 2 to initiate FSeDR reaction. Lastly, the related proteins were pulled down using streptavidin beads and probed by western blot (Figure 5A). As shown in Figure 5B, the majority of the biotinylated cellular proteins were enriched and released in lane 10, with barely left in the supernatant lane 7.

**Figure 5.**
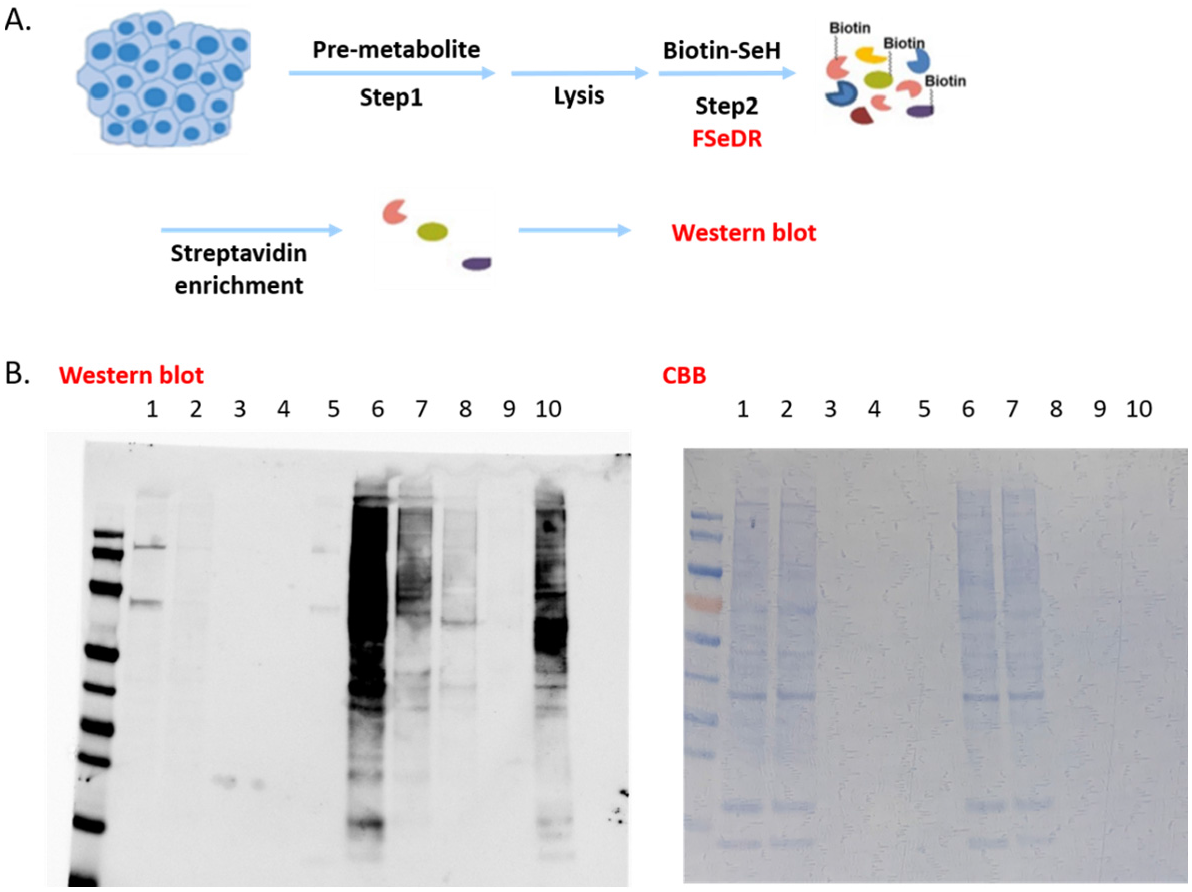
Pull down of acetylated proteins labeled by FSeDR from HEK293 cell lysates. (A) Experimental scheme for cellular pro-metabolite incorporation (step 1), protein substrates labeling by Biotin-SeH probe (step 2), enrichment with streptavidin beads, followed by western blot analysis. (B) Western blot result. Lanes 1-5 are samples from blank control; lanes 6-10 are samples from fluorine labeled cells. Briefly, lane 1 and lane 6 are input from 30% of cell lysate; lane 2 and lane 7 are the unbound elution; lane 3 and lane 8 are the first time washing elution (0.2% SDS, PBS, 4M urea); lane 4 and lane 9 are the second time washing elution (0.1% SDS, PBS); lane 5 and lane 10 are the elution of samples from the beads. Left panel: western blot with streptavidin-IRDye 680RD; right panel: CBB staining as loading control.

### Metabolic engineering of glycans using FSeDR

The success of fluorine-displacement-based labeling on acetylation compelled us to attempt it with other PTMs such as glycosylation. Thus, we explored three *N*-acetyl fluorinated monosaccharides in labeling glycosylated proteins (Figure 6A). Ac_4_ManNFAc (**23**) could hijack sialic acid biosynthetic pathway and modify silalylated glycoconjugates,^37^ while Ac_4_GalNFAc (**24**) and Ac_4_GlcNFAc (**25**) were designed to target mucin-type^38^ and O-GlcNAcylated glycoprotein,^39^ respectively. Once inside the cells, peracetylated sugars are deacetylated by cytosolic esterases, leaving the desired metabolites for further processing through the cellular machinery. We synthesized the fluorinated unnatural monosaccharides via coupling of per-acetylated amino sugars with sodium fluoroacetate. The synthesis of per-acetylated amino sugars was followed the procedure of Biswas et al ^40^ with some modifications (Scheme S3, Supporting Information). In general, the amine group was first protected as Schiff base form with 2-hydroxylnaphthaldehyde (for Man) or p-anisaldehyde (for Gal, Glc) in 1 M aqueous NaOH. Then all the four hydroxyl groups were acetylated by acetic anhydride in pyridine. After deprotection with 5 M HCl in methanol, the per-acetylated amino sugar was extracted from the reaction mixture that was quenched by aqueous Na_2_CO_3_ (Scheme S3).

**Figure 6.**
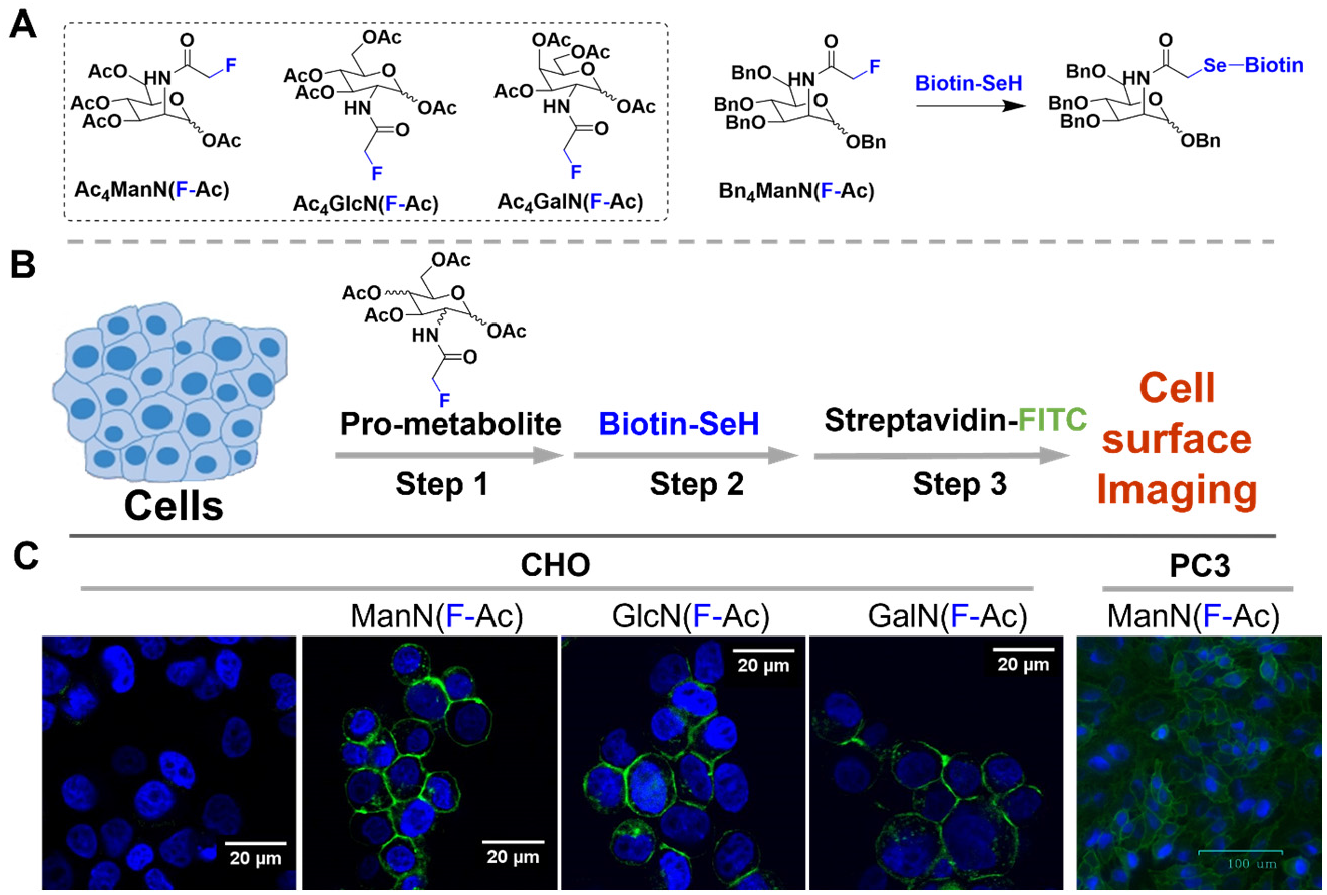
(A) N-Fluoroacetyl monosaccharide tool-kit (left) and the model reaction with Biotin-SeH probe. (B) Experimental scheme for F-glycan metabolic engineering (C) Imaging of the glycans incorporated onto cell surfaces. Nuclei (blue), FITC (green).

We suspected that after metabolic labeling, fluorine as a chemical reporter would be displayed on the cell membrane. By using FSeDR reaction, the fluorine tag could be replaced by biotin with Biotin-SeH probe. To test whether fluorinated sugars were suitable for metabolic engineering, CHO or PC-3 cells were first incubated with **23**-**25** for 36 h, then fixed with paraformaldehyde (Figure 6B). The resulted cells were treated with the Biotin-SeH (2 mM) for 8 h at 37 °C. Cell surface glycosylated proteins were finally imaged upon staining with FITC-streptavidin conjugate (Figure 6B). Compared with the cells treated with DMSO, fluorescent imaging showed intensive labelling of the cell membrane for almost all the cells samples treated with each *N*-acetyl fluorinated monosaccharide (Figure 6C). The ability to tag and convert the F-tag on PTMs other than acetylation is an important milestone in the quest for steric-free profiling of PTM substrate proteins.

### Quantitative proteomics studies of acetylation using FSeDR

With sufficient evidence to support the FSeDR based probes’ labeling, detecting, and imaging of acetylation and glycosylation, we finally decided to profile potentially new substrate proteins using acetylation as an example. Prostate cancer (PCa) is the most common male specific cancer and second leading cause of cancer-related deaths in US males. Approximately 240,000 men are diagnosed annually with PCa in the US.^41^ Almost androgen deprivation therapy is effective towards some prostate cancer patients, certain sub-cancer types such as PC3, is androgen-independent although the mechanisms are poorly understood,^42 43^ particularly in terms of the role of PTMs such as acetylation. Thus, we decide to use PC3 in this study to explore how dynamic acetylation affect prostate tumorigenesis. It is generally considered that metabolic incorporation efficiency of bioorthogonal chemical reporters are relatively low in most cell lines.^44^ Thus, to ensure the availability of fluorine labeled cofactor to acetyltransferase, fresh PC-3 cell lysates (maintaining intrinsic KATs’ activity) were directly incubated with different concentration of F-Ac-CoA, followed by western blotting to detect the F-acetylation level. In our hands, 2 mM F-Ac-CoA was found to be sufficient for the purpose (Figure 7).

**Figure 7.**
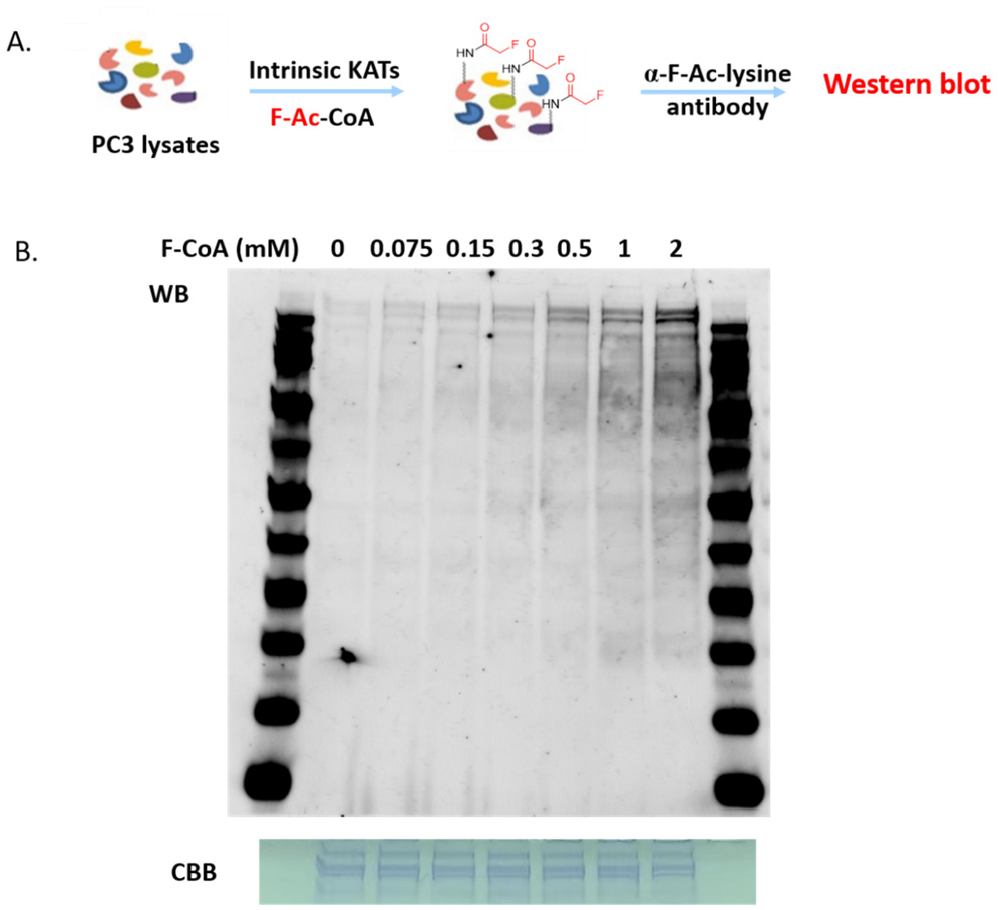
Dose-dependent F-acetylation of cell lysates *in vitro* by KATs and F-Ac-CoA. (A) Experimental scheme for direct F-acetyl incorporation in cell lysates and western blot analysis by anti-F-Acetyl lysine antibody. (B) Western blotting images of the results

MS-based quantitative proteomics has become a powerful tool in biological research.^45^ Since MS is inherently nonquantitative, the stable isotopes are introduced into the proteins, allowing relative quantification of biomolecules from different samples in the same MS analysis to rule out nonspecific background proteins.^46^ SILAC is particularly suitable for studies with extensive sample processing, such as subcellular fractionation, affinity purification of protein complex or enrichment of peptides with PTMs.^47^ Thus, we performed SILAC in tandem with FSeDR labeling to selectively enrich acetylation substrate proteins from PC3 cells cultured in heavy isotope labeled amino acid culture (Figure 8A). With a total of four independent runs, we have identified 100 enriched proteins with p value less than 0.05, among which 43 proteins have never been reported before as acetylation substrates. Preliminary validation using FSeDR labeling in cell lysates confirmed that most identified proteins are real (Figure 8B-8C). To further validate the target proteins, we picked those with significant p values, and performed knock-in amplification in HEK293 cells. As shown in Figure 8D, not only proteins such as heat shock HSPA6 and RNA helicase DDX54 have basal level of acetylation, but their acetylation levels can also be further enhanced through the addition of HADC inhibitors or the knock-in expression of the acetyltransferase GCN5.

**Figure 8.**
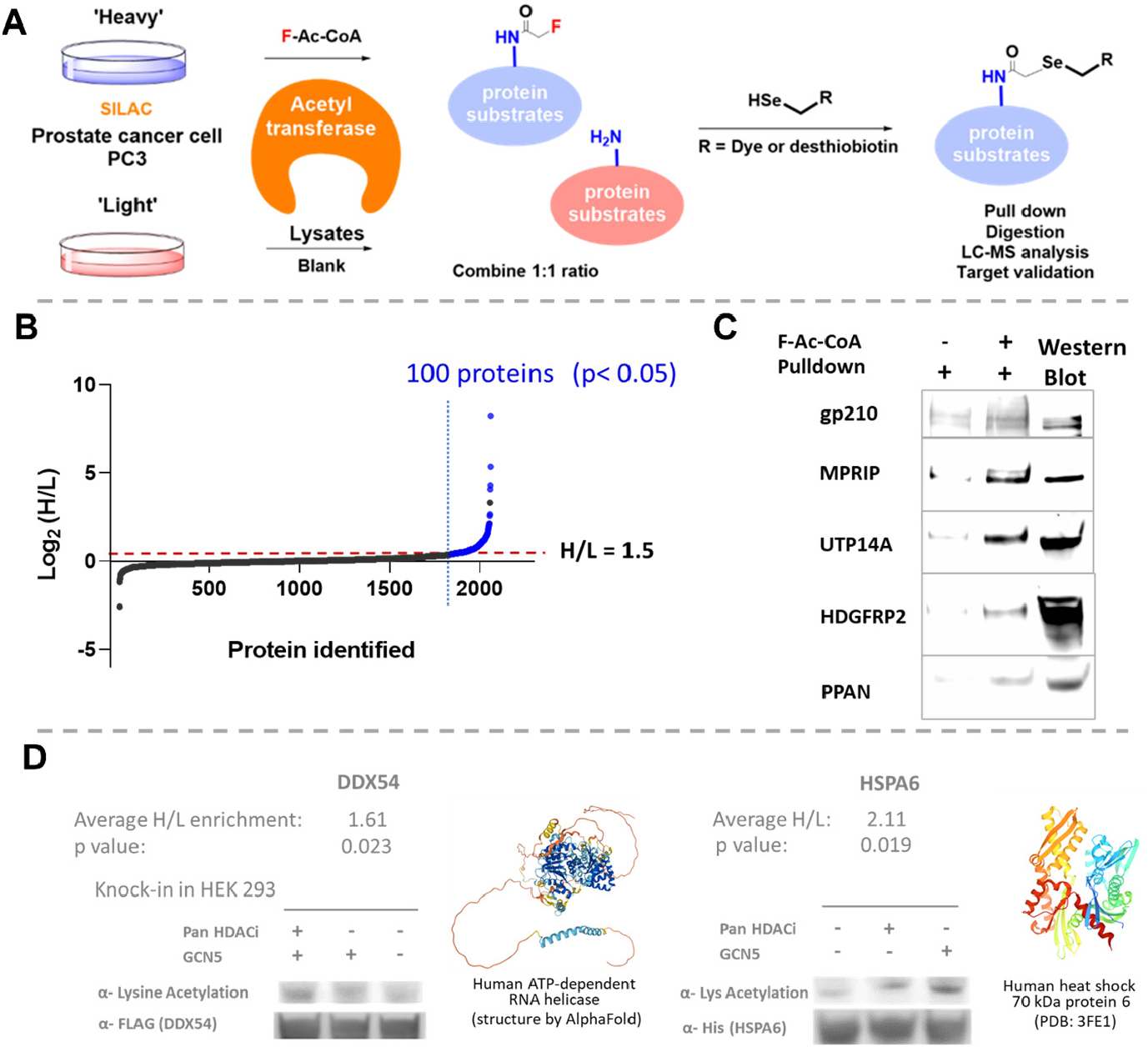
FSeDR and SILAC based chemical proteomics study of protein acetylation in model cancer cell line PC3. (A) Experimental scheme for fluorine-selenium displacement reaction in tandem with SILAC quantitative proteomics to identify acetylation substrates in target cells. (B) Proteomics results of acetylation substrates enriched in prostate cancer cells PC3. n = 4 independent repeats. (C) Validation of proteomics results in F-Ac-CoA treated cell lysates. (D) Validation of enriched protein substrates (with promising p values) using knock-in studies on HEK293 cells. GCN5: a major isoform of acetyltransferase.

## Conclusions

In summary, we have developed a fluorine-selenol displacement reaction (FSeDR) and applied it in global profiling of important PTMs such as acetylation and glycosylation. Combined with SILAC-based quantitative proteomics studies, our selenol probes revealed a good amount of novel protein substrates for acetylation. As the improved version of FTDR, FSeDR has the advantages of mild reaction condition, faster reaction kinetic, more hydrophilic conjugation product, etc. Future work would focus on proteomics studies of more disease-relevant biological systems such as the migration of cancer cell lines. With the more potent selenol probes in hand, we are also going to explore the probing of other PTM types. Nevertheless, the current proof-of-concept research has provided strong evidence that the FSeDR-based labeling strategy can be a powerful platform that allows systematic studies of a various set of PTMs and potentially many other interesting biological modifications.

## Supporting Information

The Supporting Information is available free of charge at xxxx

## Author contributions

These authors contributed equally.

## Notes

The authors declare no competing financial interests.

## Acknowledgments

This work has been supported by NIH grant 5R35GM133468-03 and Temple University Startup Funding. Rongsheng E. Wang is a Cottrell Scholar of Research Corporation for Science Advancement. Support for the Wistar Proteomics and Metabolomics Facility was provided by P30 CA010815 and S10 OD023586. Support for the NMR facility at Temple University by a CURE grant from the Pennsylvania Department of Health is gratefully acknowledged.

## References

1. Ponomarenko, E. A.; Poverennaya, E. V.; Ilgisonis, E. V.; Pyatnitskiy, M. A.; Kopylov, A. T.; Zgoda, V. G.; Lisitsa, A. V.; Archakov, A. I., The Size of the Human Proteome: The Width and Depth. Int J Anal Chem. 2016, 2016, 7436849.

2. Buuh, Z. Y.; Lyu, Z.; Wang, R. E., Interrogating the roles of post-translational modifications of Non-histone proteins. J. Med. Chem. 2018, 61, 3239–3252.

3. Ali, I.; Conrad, R. J.; Verdin, E.; Ott, M., Lysine Acetylation Goes Global: From Epigenetics to Metabolism and Therapeutics. Chem. Rev. 2018, 118 (3), 1216–1252.

4. Marmorstein, R.; Zhou, M.-M., Writers and Readers of Histone Acetylation: Structure, Mechanism, and Inhibition. Cold Spring Harb Perspect Biol. 2014, 6 (7), a018762.

5. Dejgaard, S.; Nicolay, J.; Taheri, M.; Thomas, D. Y.; Bergeron, J. J. M., The ER glycoprotein quality control system. Curr Issues Mol Biol. 2004, 6, 29–42.

6. Schjoldager, K. T.; Narimatsu, Y.; Joshi, H. J.; Clausen, H., Global view of human protein glycosylation pathways and functions. Nat. Rev. Mol. Cell Biol. 2020, 21, 729–749.

7. Varki, A., Evolutionary Forces Shaping the Golgi Glycosylation Machinery: Why Cell Surface Glycans Are Universal to Living Cells. Cold Spring Harb Perspect Biol. 2011, 3, a005462.

8. Taniguchi, N.; Honke, K.; Fukuda, M.; Narimatsu, H.; Yamaguchi, Y.; Angata, T., Handbook of Glycosyltransferases and Related Genes. 2014.

9. Varki, A.; Cummings, R. D.; Esko, J. D.; Stanley, P.; Hart, G. W.; Aebi, M.; Darvill, A. G.; Kinoshita, T.; Packer, N. H.; Prestegard, J. H.; Schnaar, R. L.; Seeberger, P. H., Essentials of Glycobiology, 3rd edition. Cold Spring Harbor Laboratory Press 2017.

10. Munkley, J.; Scott, E., Targeting Aberrant Sialylation to Treat Cancer. Medicines (Basel). 2019, 6 (4), 102.

11. Pinho, S. S.; Reis, C. A., Glycosylation in cancer: mechanisms and clinical implications. Nature Reviews Cancer 2015, 15, 540–555.

12. Goldin, A.; Beckman, J. A.; Schmidt, A. M.; Creager, M. A., Advanced Glycation End Products. Circulation 2006, 114, 597–605.

13. Narita, T.; Weinert, B. T.; Choudhary, C., Functions and mechanisms of non-histone protein acetylation. Nat. Rev. Mol. Cell Biol. 2019, 20, 156–174.

14. ChanKim, S.; Sprung, R.; Chen, Y.; Xu, Y.; Ball, H.; Pei, J.; Cheng, T.; Kho, Y.; Xiao, H.; Xiao, L.; Grishin, N. V.; White, M.; Yang, X.-J.; Zhao, Y., Substrate and Functional Diversity of Lysine Acetylation Revealed by a Proteomics Survey. Mol. Cell 2006, 23 (4), 607–618.

15. Choudhary, C.; Kumar, C.; Gnad, F.; Nielsen, M. L.; Rehman, M.; Walther, T. C.; Olsen, J. V.; Mann, M., Lysine Acetylation Targets Protein Complexes and Co-Regulates Major Cellular Functions. Science 2009, 325 (5942), 834–840.

16. Kaiser, C.; James, S. R., Acetylation of insulin receptor substrate-1 is permissive for tyrosine phosphorylation. BMC Biology 2004, 2, 23.

17. Min, S.-W.; Cho, S.-H.; Zhou, Y.; Schroeder, S.; Haroutunian, V.; Seeley, W. W.; Huang, E. J.; Shen, Y.; Masliah, E.; Mukherjee, C.; Meyers, D.; Cole, P. A.; Ott, M.; Gan, L., Acetylation of Tau Inhibits Its Degradation and Contributes to Tauopathy. Neuron 2010, 67 (6), 953–966.

18. Wan, J.; Zhan, J.; Li, S.; Ma, J.; Xu, W.; Liu, C.; Xue, X.; Xie, Y.; Fang, W.; Chin, Y. E.; Zhang, H., PCAF-primed EZH2 acetylation regulates its stability and promotes lung adenocarcinoma progression. Nucleic Acids Res. 2015, 43 (7), 3591–3604.

19. Li, L.; Yang, X.-J., Tubulin acetylation: responsible enzymes, biological functions and human diseases. Cell Mol Life Sci. 2015, 72, 4237–4255.

20. AdrianDrazic; M. Myklebust, L.; Rasmus Ree; Thomas Arnesen, The world of protein acetylation. Biochimica et Biophysica Acta (BBA) - Proteins and Proteomics 2016, 1864 (10), 1372–1401.

21. Lyu, Z.; Zhao, Y.; Buuh, Z. Y.; Gorman, N.; Goldman, A. R.; Islam, M. S.; Tang, H.-Y.; Wang, R. E., Steric-Free Bioorthogonal Labeling of Acetylation Substrates Based on a Fluorine-Thiol Displacement Reaction. J. Am. Chem. Soc. 2021, 143 (3).

22. Li1, Y.; Silva, J. C.; Skinner, M. E.; Lombard, D. B., Mass spectrometry-based detection of protein acetylation. Methods Mol Biol. 2013, 1077, 81–104.

23. Shaw, P. G.; Chaerkady, R.; Zhang, Z.; Davidson, N. E.; Pandey, A., Monoclonal Antibody Cocktail as an Enrichment Tool for Acetylome Analysis. Anal. Chem. 2011, 83 (10), 3623–3626.

24. Hendrickson, O. D.; Zherdev, A. V., Analytical Application of Lectins. Critical Reviews in Analytical Chemistry 2018, 48 (4), 279–292.

25. Schjoldager, K. T.; Narimatsu, Y.; Joshi, H. J.; Clausen, H., Global view of human protein glycosylation pathways and functions. Nat. Rev. Mol. Cell Biol 2020, 21, 729–749.

26. Palaniappan, K. K.; Bertozzi, C. R., Chemical glycoproteomics. Chem. Rev. 2016, 116, 14277–14306.

27. Song, J.; Zheng, Y. G., Bioorthogonal Reporters for Detecting and Profiling Protein Acetylation and Acylation. SLAS discovery 2019, 25 (2), 148–162.

28. S. McKay, C.; M.G. Finn, Click Chemistry in Complex Mixtures: Bioorthogonal Bioconjugation. Chemistry&Biology 2014, 21 (9), 1075–1101.

29. Wratil, P. R.; Horstkorte, R.; Reutter, W., Metabolic glycoengineering with N-acyl side chain modified mannosamines. Angew. Chem. Int. Ed. 2016, 55, 9482–9512.

30. Yang, C.; Mi, J.; Feng, Y.; Ngo, L.; Gao, T.; Yan, L.; Zheng, Y. G., Labeling Lysine Acetyltransferase Substrates with Engineered Enzymes and Functionalized Cofactor Surrogates. J. Am. Chem. Soc. 2013, 135 (21), 7791–7794.

31. Xiong, D.-C.; Zhu, J.; Han, M.-J.; Luo, H.-X.; Wang, C.; Yu, Y.; Ye, Y.; Tai, G.; Ye, X.-S., Rapid probing of sialylated glycoproteins in vitro and in vivo via metabolic oligosaccharide engineering of a minimal cyclopropene reporter. Org. Biomol. Chem 2015, 13, 3911–3917.

32. Han, Z.; Chou, C.-w.; Yang, X.; Bartlett, M. G.; Zheng, Y. G., Profiling Cellular Substrates of Lysine Acetyltransferases GCN5 and p300 with Orthogonal Labeling and Click Chemistry. ACS Chem. Biol. 2017, 12 (6), 1547–1555.

33. He, M.; Han, Z.; Liu, L.; Zheng, Y. G., Chemical Biology Approaches for Investigating the Functions of Lysine Acetyltransferases. Angew. Chem. Int. Ed. Engl. 2017, 57, 1162–1184.

34. Reich, H. J.; Hondal, R. J., Why nature chose selenium. ACS Chem. Biol. 2016, 11, 821–841.

35. Pálla, T.; Mirzahosseini, A.; Noszál, B., Species-Specific, pH-Independent, Standard Redox Potential of Selenocysteine and Selenocysteamine. Antioxidants 2020, 9, 465.

36. Dery, S.; Reddy, P. S.; Dery, L.; Mousa, R.; Dardashti, R. N.; Metanis, N., Insights into the deselenization of selenocysteine into alanine and serine. Chem. Sci. 2015, 6, 6207–6212.

37. Du, J.; Meledeo, M. A.; Wang, Z.; Khanna, H. S.; Paruchuri, V. D. P.; Yarema, K. J., Metabolic glycoengineering: Sialic acid and beyond. Glycobiology 2009, 19, 1382–1401.

38. Hang, H. C.; Yu, C.; Kato, D. L.; Bertozzi, C. R., A metabolic labeling approach toward proteomic analysis of mucin-type O-linked glycosylation. Proc Natl Acad Sci U S A. 2003, 100 (25), 14846–14851.

39. Zaro, B. W.; Yang, Y.-Y.; Hang, H. C.; Pratt, M. R., Chemical reporters for fluorescent detection and identification of O-GlcNAc-modified proteins reveal glycosylation of the ubiquitin ligase NEDD4-1. Proc. Natl. Acad. Sci. U.S.A 2011, 108 (20), 8146–8151.

40. Biswas, N. N.; Yu, T. T.; Kimyon, Ö.; Nizalapur, S.; Gardner, C. R.; Manefield, M.; Griffith, R.; Black, D. S.; Kumar, N., Synthesis of antimicrobial glucosamides as bacterial quorum sensing mechanism inhibitors. Bioorg. Med. Chem. 2017, 25, 1183–1194.

41. Siegel, R.; Naishadham, D.; Jemal, A., Cancer statistics, 2013. CA Cancer J Clin. 2013, 63 (1), 11–30.

42. Nelson, W. G.; Marzo, A. M. D.; Isaacs, W. B., Prostate Cancer. N Engl J Med. 2003, 349, 366–381.

43. Saraon, P.; Drabovich, A. P.; Jarvi, K. A.; Diamandis, E. P., Mechanisms of Androgen-Independent Prostate Cancer. EJIFCC. 2014, 25 (1), 42–54.

44. D., P. N.; Fermaintt, C. S.; Rodriguez, A. C.; McCombs, J. E.; Nischan, N.; Kohler., J. J., Cellular metabolism of unnatural sialic acid precursors. Glycoconj. J. 2015, 32, 515–529.

45. Bantscheff, M.; Schirle, M.; Sweetman, G.; Rick, J.; Kuster, B., Quantitative mass spectrometry in proteomics: a critical review. Anal. Bioanal. Chem. 2007, 389, 1017–1031.

46. Schubert, O. T.; Röst, H. L.; Collins, B. C.; Rosenberger, G.; Aebersold, R., Quantitative proteomics: challenges and opportunities in basic and applied research. Nat. Protoc. 2017, 12, 1289–1294.

47. Chen, X.; Wei, S.; Ji, Y.; Guo, X.; Yang, F., Quantitative proteomics using SILAC: Principles, applications, and developments. Proteomics 2015, 15, 3175–3192.

