## Supporting Information for "Steric-free bioorthogonal profiling of cellular acetylation and glycosylation via a fluorine-selenol displacement reaction (FSeDR)"

#### Table of Contents

Materials and Methods

Chemical Synthesis ……………………………………………………………………S2

Biological Experiments ………………………………………………………………..S6

Compound Characterizations ……………………………………………………………S9

Supplementary Figures …………………………………………………………………S16

### Materials and Methods

#### *Chemical Synthesis*

##### General Information:

Chemical reagents and solvents were purchased from commercial resources such as VWR, Thermo Fisher, and Sigma Aldrich, and were used directly without further purification. Analytical TLC was carried out with Silica Gel 60 $F_{254}$ plates (EMD Chemicals). The chemicals on TLC were either visualized by UV 254 nm (UV lamp, Chemglass Life Sciences) or stained by phosphomolybdic acid or $KMnO_4$ oxidation. Compound purification was performed by normal-phase flash column chromatography on columns manually loaded with silica gel grade 60 (230-400 mesh, Fisher Scientific) or by reverse-phase Combi-Flash on prepacked C18 columns (Teledyne ISCO). Further purification by preparative high-performance liquid chromatography (HPLC) was implemented on Waters 1525 series that consist of a 2489 UV/vis detector, 1525 binary pump, and an XBridge Prep C18 column. Routine mass spectrometry analysis was done using liquid chromatography-mass spectrometry (LC-MS) Agilent series. High resolution LC-MS analysis was performed on an Agilent 6520 Accurate-Mass Quadrupole-Time-of-Flight (Q-TOF) coupled with an electrospray ionization source. For NMR analysis, $^{1}$H NMR and $^{13}$C NMR spectra were recorded on 400 MHz or 500 MHz Bruker Advance. The raw data were processed with MestReNova, and the chemical shifts were reported in parts per million (ppm) downfield from the internal standard tetramethylsilane (TMS).

**Scheme S1**. Synthesis of selenol probes.

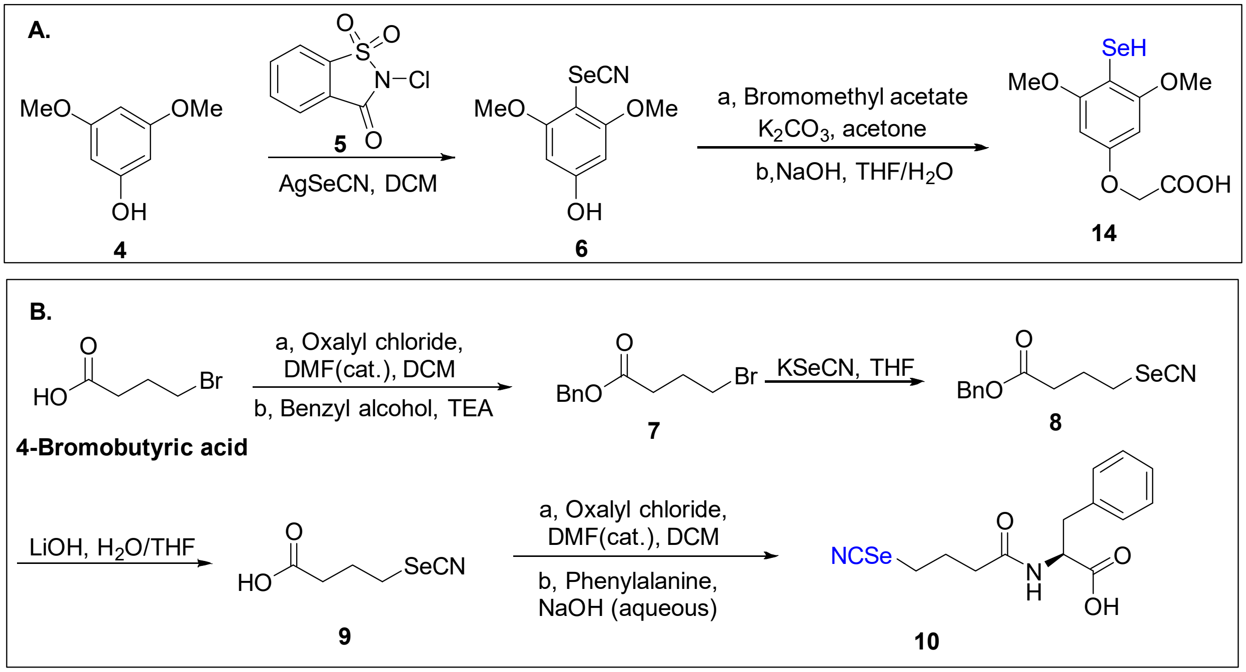

**Scheme S2.** Synthesis of desthiobiotin-SeCN probe.

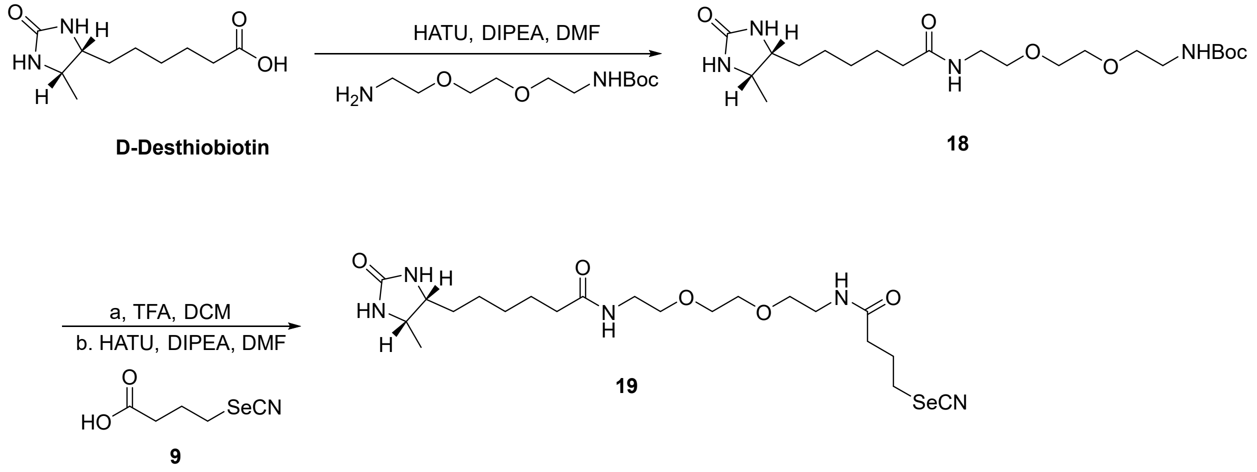


#### General Procedure for Comparison of Selenol/Thiol Probes:

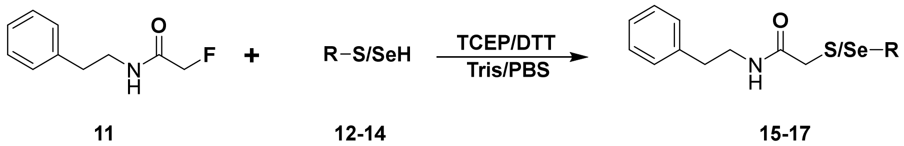


**FTDR**: 2 µL substrate **11** (200 mM stock in $CH_3CN$) was mixed with 5 µL model thiol phenol probe **12** (100 mM stock in $CH_3CN$) and 2 µL TCEP (1 M stock in $H_2O$, pH 7.2). Tris buffer was added to make a total volume of 20 µL and NaOH (500 mM) was added to adjust the final pH value to 8.5.

**FSeDR:** 5 µL model aliphatic selenol probe **13** (100 mM stock in $CH_3CN$) or model selenol phenol probe **14** (100 mM stock in $CH_3CN$) was first treated with 2 µL DTT (1 M stock in $H_2O$) at room temperature for 10 min. Then 2 µL substrate **11** (200 mM stock in $CH_3CN$) was added. DPBS buffer was added to make a total volume of 20 µL and NaOH (50 mM) was added to adjust the final pH value to 7.4.

The mixture was reacted at 37 °C. Approximately 5 µL of the reaction mixture was taken out at 4 h and was mixed with 20 µL 0.1% TFA/ACN to quench the reaction. The samples were analyzed by LC-MS.

#### General Procedure for Testing Cyano-Deprotection Conditions:

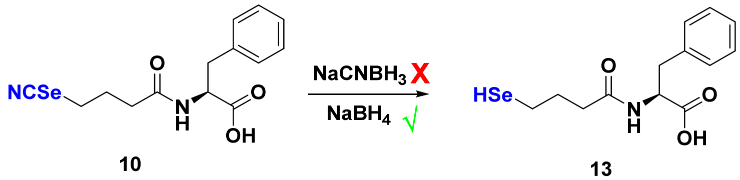


5 µL cyano protected model aliphatic selenol probe **10** (100 mM stock in $CH_3CN$) was mixed with 5 µL $NaCNBH_3$ or $NaBH_4$ (fresh 500 mM stock in $H_2O$) at room temperature for 10 min. Then 5

μL of the reaction mixture was taken out and was mixed with 20 μL 0.1% TFA/ACN to quench the reaction. The samples were analyzed by LC-MS. Probe **10** was found to be fully converted to probe **13** by treating $NaBH_4$ (Figure 3.3A-C) while no reaction was found by treating $NaCNBH_3$. Besides, no reaction was detected still after incubating **10** with $NaCNBH_3$ for 2 h or with double amount of $NaCNBH_3$ for 2 h.

**General Procedure for Testing the Reduction of Selenium Dimers:**

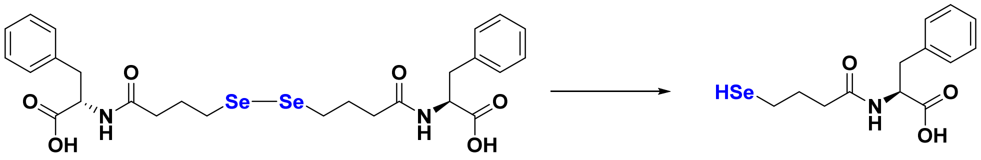


**13-dimer** **13**

2 μL **13-dimer** (20 mM stock in $H_2O$) was mixed with I. 2 μL TCEP (500 mM stock in $H_2O$); II. 2 μL DTT (500 mM stock in $H_2O$); III. 2 μL $NaBH_4$ (fresh 100 mM stock in $H_2O$); IV. 2 μL $NaCNBH_3$ (fresh 100 mM stock in $H_2O$) at the room temperature. DPBS was added to adjust the final volume to 20 μL and adjust the final pH value to 7.4. After 1 h incubation, all the samples were analyzed by LC-MS.

**Optimizing Conditions - pH Titration of the fluorine-selenol displacement reaction:**

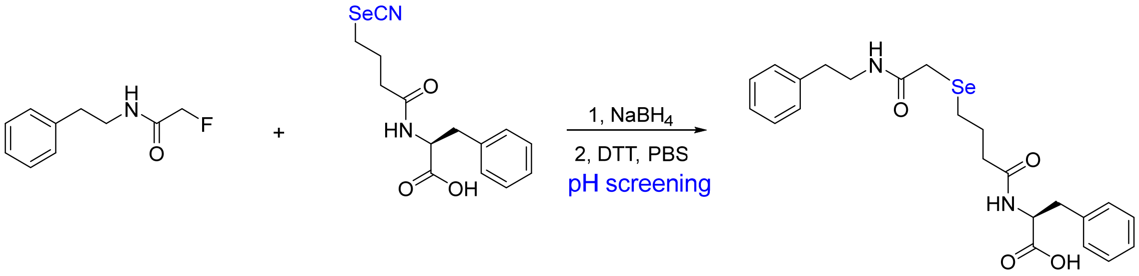


**11** **13** **16**

5 μL probe **10** (200 mM stock in $CH_3CN$) was mixed with 4 μL $NaBH_4$ (1 M stock in $H_2O$) at room temperature for 10 min. Then 4 μL DTT (1 M stock in $H_2O$) and 2 μL substrate **11** (200 mM stock in $CH_3CN$) was added. DPBS buffer was added to make a total volume of 20 μL and HCl (1 M) or NaOH (500 mM) was added to adjust the final pH value to 6.0, 7.2, 8.0. Thus, the final concentration of substrate **11**, selenol probe, and DTT was 20 mM, 50 mM, and 200 mM, respectively. The mixture was reacted at 37 °C. Approximately 5 μL of the reaction mixture was taken out at 5 h and was mixed with 20 μL 0.1% TFA/ACN to quench the reaction. The samples were analyzed by LC-MS.

**Scheme S3.** Synthesis of N-Acetyl fluorinated monosaccharides.

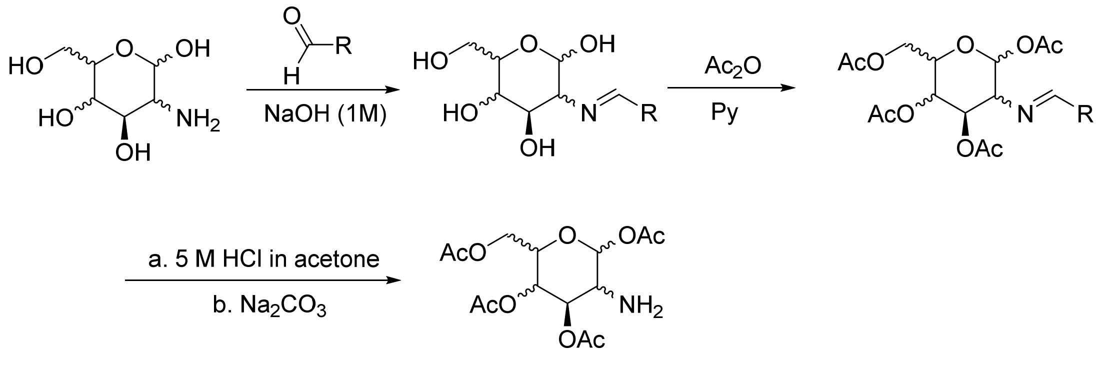


**20 - 22**

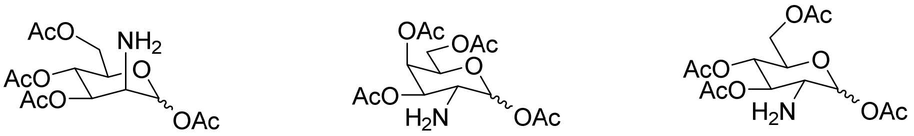

**Ac$_4$ManNH$_2$**
**20**

**Ac$_4$GalNH$_2$**
**21**

**Ac$_4$GlcNH$_2$**
**22**

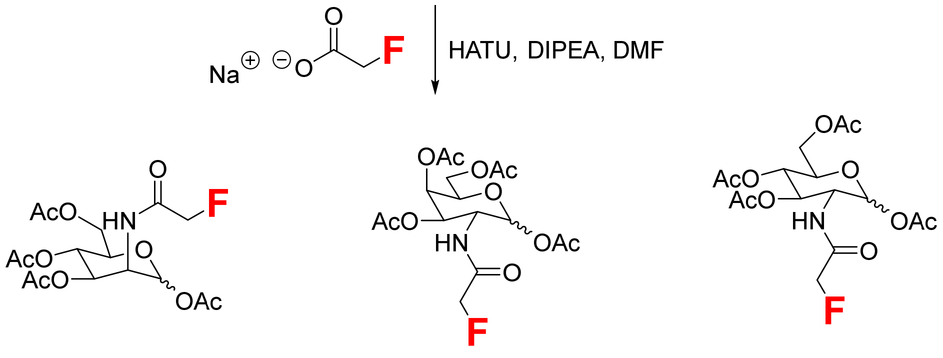


**Ac$_4$ManNFAc**
**23**

**Ac$_4$GalNFAc**
**24**

**Ac$_4$GlcNFAc**
**25**

**General Procedure for Synthesis of Acetated Monosaccharide Intermediates 20 - 22:** Mannosamine/galacosamine/glucosamine hydrochloride (5 mmol, 1 equiv) was dissolved in 1 M NaOH (5 mL) solution at 0 °C and 2-hydroxylnaphthaldehyde (for Man, 5.5 mmol, 1.1 equiv) or p-anisaldehyde (for Gal, Glc, 5.5 mmol, 1.1 equiv) was added dropwise. The reaction was then warmed to room temperature and stirred overnight. The precipitated white solid was collected by vacuum filtration, washed with cold water, ethanol and diethyl ether, and then dried under vacuum to give the imine intermediate as a white solid. The intermediated was dissolved in dry pyridine (5 mL) and acetic anhydride (25 mmol, 5 equiv) was then added dropwisely. After stirred at room temperature overnight, the reaction was quenched with water and ethyl acetate. The organic layer was washed with water twice and dried over anhydrous sodium sulfate. After concentrated under vacuum, the resulted white solid was dissolved in acetone (5 mL) and treated with HCl (5 M, 2 mL). The solution was stirred for 1 h before saturated sodium bicarbonate was added. The product was then extracted with ethyl acetate, dried over sodium sulfate, and concentrated under vacuum.

The crude mixture was purified via flash column chromatography (DCM/$CH_3OH$: 30/1) to afford the final product.

**General Procedure for Synthesis of N-Acetyl F-Monosaccharides 23 - 25:**
Sodium fluoroacetate (0.37 mmol, 1.5 equiv), HATU (0.37 mmol, 1.5 equiv) and DIPEA (0.50 mmol, 2 equiv) were mixed in dry DMF (3 mL) for 20 min. Then the amine intermediate **1-3** (0.25 mmol, 1 equiv) dissolved in DMF (1 mL) was added. After stirred at room temperature for 4 h, the reaction mixture was quenched with water and the product was extracted with ethyl acetate. The organic layer was washed with water twice, dried over anhydrous sodium sulfate, concentrated under vacuum. The crude mixture was purified via flash column chromatography (Hexane/ Ethyl acetate: 2/ 1) to afford the final product.

#### *Biological Experiments*

**Cell Culturing**
Jurkat, CHO, MCF-7 cells were cultured in RPMI-1640 medium. Hela and HEK293 cells were cultured in DMEM medium. PC-3 cells were cultured in DME/F-12 1:1 medium. All medium containing 10% FBS, 100 IU/mL penicillin, and 100 μg/mL streptomycin. All cells were incubated in a humidified incubator at 37 °C with 5% $CO_2$.

**Fluorination and Biotinylation of Bovine Serum Albumin (BSA)**
400 μL of BSA (2.4 mg/mL in DPBS, pH 7.6) was first reacted with 14.8 μL of NHS-fluoroacetate (20 mM stock in DMSO) at room temperature for 2 h. The fluorinated BSA was purified by cold methanol precipitation to remove unconjugated linkers and then the pellet was redissolved in 300 μL water (2.6 mg/mL, ~39 μM). For biotinylation, when using FTDR labeling, 8 μL fluorinated BSA (39 μM stock in water) was mixed with 4 μL TCEP (100 mM stock in water), 2 μL Biotin-SH (30 mM stock in DMSO), 1 μL Tris buffer (1 M stock in water) and 5 μL NaOH (200 mM stock in water). The final reaction volume was 20 μL and pH was 8.5; when FSeDR was used, 2 μL Biotin-SeCN probe **20** (30 mM stock in $CH_3CN$) was mixed with 1 μL $NaBH_4$ (300 mM stock in water) at the room temperature for 10 min. Then 4 μL DTT (100 mM stock in water), 5 μL PBS (250 mM stock in water) and 8 μL fluorinated BSA (39 μM stock in water) were added to the activated probe solution. The final reaction volume was 20 μL and pH was 7.2. Both of reaction mixture were incubated at 37 °C for the indicated time point (3 h, 6 h). 6 μL samples from each time point were loaded onto 8% Bis-Tris SDS-PAGE and transferred onto PVDF membranes. The blot was blocked with biotin-free casein buffer (Sigma Aldrich) for 1 h, stained with streptavidin-IRDye 680RD (LI-COR) at a dilution of 1/2000 for 1 h, followed by 3 times washings with PBST (containing 0.1% Tween-20). The biotinylated proteins were detected by the LI-COR Odyssey Fc Imaging System (700 nm channel, Ex 685 nm/Em 730 nm). At the end, the blot was stained with CBB to show equal loading amount for each sample.

**FSeDR-Based Pull Down of Protein Substrates Followed by Western Blot Analysis**

- **BSA model**

40 μL Biotin-SeCN probe **20** (30 mM stock in $CH_3CN$) was mixed with 20 μL $NaBH_4$ (300 mM stock in water) at the room temperature for 10 min. Then 80 μL DTT (100 mM stock in water),

100 μL PBS (250 mM stock in water) and 160 μL fluorinated BSA (39 μM stock in water) were added to the activated probe solution. The final reaction volume was 400 μL and pH was 7.2. After incubated at 37 °C for 6 h, the reaction was added with 1.5 mL cold methanol to precipitate proteins and remove extra linkers. The white pellet was redissolved in 100 μL DPBS and the biotinylated BSA concentration was determined to be 2.4 mg/mL by nanodrop analysis. For sample 1, 50 μL biotinylated BSA in DPBS (1 mg/mL) was mixed with 50 μL streptavidin magnetic beads (New England BioLabs) at room temperature for 1 h. The beads were washed with PBS three times. For sample 2 and 3, 0.1% and 0.2% SDS was added to the input buffer and washing buffer separately while other conditions stayed same. Then the beads were added with gel loading buffer and heated at 95 °C for 10 min to elute enriched proteins. 10% of input proteins, unbounded supernatant, washes, and elution were loaded to 8% SDS-PAGE gel. After transferred to PVDF membrane, the blot was blocked and analyzed as the procedure mentioned before. The best buffer for dissolving input proteins was found to be PBS with 0.1% SDS. For washing buffer, DPBS containing up to 0.2% SDS would not disrupt streptavidin biotin interaction.

- **HEK293 cell lysates**

HEK293 cells at 80% confluency were treated with ethyl fluoroacetate (2 mM) for 6 h and then lysed with CelLytic M buffer (Sigma-Aldrich) containing Protease Inhibitor Cocktail (cOmplete, EDTA-free, Roche). The lysates were purified by methanol precipitation and the precipitated proteins were redissolved in DPBS containing 1% SDS and the resulting proteome (~ 2 mg/mL, pH 7.2) was added with 50 mM DTT, 3 mM sodium borohydride activated Biotin-SeH probe (compound **10** → **13**). After incubated at 37°C for 8 h, the extra probes in the reaction mixture were removed by methanol precipitation. The protein pellet was redissovled in PBS buffer (0.1% SDS, pH 7.2). 50 μL biotinylated proteome in DPBS (~ 1 mg/mL) was mixed with 50 μL streptavidin magnetic beads (New England BioLabs) at room temperature for 1 h. After removing unbound lysates, the beads were washed with 0.2% SDS in DPBS containing 4 M urea once and 0.1% SDS in DPBS twice. Then enriched biotinylated proteins were eluted by heating in the loading buffer. 30% of input proteins, unbounded supernatant, washes, and elution were loaded to 4-12% Bis-Tris SDS-PAGE gel. Distribution of biotinylated proteins in the pull-down process were analyzed by western blotting using streptavidin IRDye 680RD as mentioned above.

**Labelling of Cell-Surface using Fluorine Modified Sugars**

Suspension Jurkat cells were seeded in 24-well culture plates at a density of $0.2 \times 10^6$ cells/mL, supplied with 150 μM DMSO or fluorine modified sugars including $Ac_4$ManNFAc (**4**), $Ac_4$GalNFAc (**5**), and $Ac_4$GlcNFAc (**6**) at indicated concentration (100, 150, 200 μM) in 1 mL of RPMI medium. After 2 days incubation, the cells were harvested with 1.5 mL Eppendorf tubes, pelleted (2000 rpm, 5 min), washed with 1 mL PBS twice. Then the cells were transferred into round bottom 96 well plate and fixed in 3.2% paraformaldehyde in PBS solution (200 μL) for 20 min. For adherent cells, the cells were seeded in flat bottom 96-well culture plates at a density of $0.01 \times 10^6$ cells/mL. Next day, the cells were supplied with DMSO or fluorine modified sugars including $Ac_4$ManNFAc (**4**), $Ac_4$GalNFAc (**5**), and $Ac_4$GlcNFAc (**6**) at indicated concentration (100, 150, 200 μM) in 100 μL of correspond medium. After incubation for indicated times (6, 12, 24, 48 h), the cells were rinsed with PBS twice, fixed in 3.2% paraformaldehyde in PBS solution (100 μL) for 20 min.

After another round of rinses, cells were incubated in reducing solution (100 µL, 50 mM DTT, PBS, pH 7.6) for 30 min. At this point, the cells became ready for labeling by biotin probes. The reduced cells were then labelled with Biotin-SeH (80 µL; for Biotin-SeCN probe **10**, 3 mM probe **10** was first deprotected with 15 mM $NaBH_4$ for 10 min, then 50 mM DTT was added; for Biotin-SeH dimer probe **18**, 3 mM **18** with 75 mM DTT were used. PBS, pH 7.2) for 8 h at 37 °C, followed by washing with reducing solution once and PBS twice. The cells were blocked with 1% BSA in PBS for 1 h and then treated with FITC streptavidin (BD Phamingen, 2 µg /mL) in 1% BSA, PBS for 30 min at room temperature. After washed in PBS three times, the Jurkat cells were suspend in 1% BSA PBS solution and analyzed by flow cytometry (BD Accuri C6, 20000 cells were counted). For adherent cells, cells were stained with Hoechst 33342 (1 µg/mL) for 10 min and washed with PBS three times. Cellular fluorescence was examined using the ZOE fluorescent microscope imager (Bio-Rad Laboratories).

**Dose-dependent F-acetylation of cell lysates *in vitro* by KATs and F-Ac-CoA**

PC-3 cells were cultured in DME/F-12 1:1 medium (HyClone) containing 10% FBS, 100 IU/mL penicillin, and 100 µg/mL streptomycin and incubated in a humidified incubator at 37 °C with 5% $CO_2$. The cells were lysed with M-PER Mammalian Protein Extraction Reagent (ThermoFisher) at 80% confluency. Protein concentration was determined by the BCA assay and lysates (1 mg/mL) was incubated with different concentrations of F-Ac-CoA (0, 0.075, 0.15, 0.3, 0.5, 1, 2 mM) for 4 h at 30 °C. The F-acetylation level of each lysates sample were evaluated by western blot using the anti-acetylated-lysine MultiMab™ antibody (Ac-$K^2$-100, Cell Signaling, a mixture of monoclonal antibodies that can recognize acetyl-lysine and F-acetyl lysine).

**FSeDR based SILAC analysis of acetylated proteins in PC-3 cells**

PC-3 cells were cultured in DMEM:F-12 for SILAC medium (Fisher 88370) containing 10% dialyzed FBS (R&D S12850H), 100 IU/mL penicillin, 100 µg/mL streptomycin, and 17.25 mg/L L-proline (Fisher 88211, to prevent Arg to Pro conversion). For light media, 91.25 mg/L L-Lysine-2HCl (Fisher 88429) and 147.5 mg/L L-Arginine-HCl (Fisher 88427) were added; for heavy media, 113.2 mg/L L-Lysine-2HCl, $^{13}C_6$, $^{15}N_2$ (Cambridge CNLM-291-H-PK) and 154.2 mg/L L-Arginine-HCl, $^{13}C_6$, $^{15}N_4$ (Fisher 89990) were added. 100,000 cells were seeded in a 100 mm cell culture dish in 10 mL culture media (light/heavy). After 8 doublings, cells at ~80% confluency was scraped and lysed in M-PER™ Mammalian Protein Extraction Reagent with 1x EDTA-free protease inhibitor cocktail (Roche). Lysates was cleared by centrifugation at 14,000 ×g, 4 °C for 5 min. Protein concentration of the supernatant was determined by a BCA assay. For maximum depth of analysis and a possible repeat, no less than 250 µg protein per condition was prepared. Lysates (~1 mg/mL) from cells cultured in light media were incubated with 2 mM F-Ac-CoA at 30 °C for 4 h while lysates from heavy media cultured cells were incubated with same volume of water as a control. Then the proteins in lysates were purified by cold methanol precipitation. The white pellets were redissolved in DPBS containing 1% SDS and the resulting proteomes (~ 2 mg/mL, pH 7.2) were added with 3 mM Desthiobiotin-SeCN probe **23**, 15 mM $NaBH_4$, and 50 mM DTT. After incubated at 37 °C for 8 h, the proteomes were purified by methanol precipitation. The collected protein pellets were redissovled in DPBS buffer (0.1% SDS, pH 7.2) and resulted proteome solutions were mixed with 0.6 mL streptavidin magnetic beads for 2 h at the room temperature. After removing supernatant, the beads were washed with DPBS containing 0.2%

SDS, 4 M urea once and 0.1% SDS in DPBS twice. The desthiobiotinylated proteins were released by incubating beads with 5 mM competition reagent Biotin solution at 37 °C for 15 min, three times. The elution solutions were combined and lyophilized. The resulted white powders were redissolved in 20 μL Tris buffer containing 8 M urea. Then 10 mM DTT was added and incubated for 20 min at 30 °C followed by incubation with 20 mM iodoacetamide in dark for 45 min at the room temperature. The alkylation step was quenched with 20 mM DTT in dark for 30 min. After diluted with 100 mM Tris buffer (pH 8.0) (urea final concentration to 4 M), the proteins were digested with trypsin protease (1:100 w/w) for 4 h at 37 °C. Next, another round of digestion was performed overnight after further dilute protein solution with 100 mM Tris buffer to achieve 2 M urea. The resulted peptide solution was cleared by centrifuge at 2500 g for 5 min and purified by C18 columns (The Nest Group). The elution solutions were combined, lyophilized, and redissolved in pure water. The tryptic digests were analyzed using a standard 95 min run on a Thermo Q Exactive Plus mass spectrometer. MS data were searched with full tryptic specificity against the UniProt human proteome database (10/02/2020).

**Compound Characterization:**

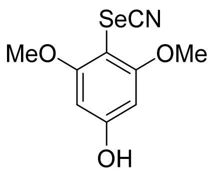


**6**

To a solution of N-chlorosaccharin (959 mg, 4.4 mmol) in dichloromethane (20 mL) was added silver selenocyanate (937 m g, 4.4 mmol). A white solid crashed out upon addition and the reaction mixture was kept stirring for 1 h. 3,5-dimethoxyphenol (616 mg ,4 mmol) was then added, and the reaction mixture was stirred for another 2 h. Then the reaction mixture was vacuum filtered, and the filtrate was vacuum concentrated. The crude mixture was purified via flash column chromatography (hexane/ethyl acetate: 2/1) to afford compound **6** as an orange solid (150 mg, 15% yield). $^{1}$H NMR (500 MHz, $CDCl_3$): $\delta$ 6.11 (s, 2H), 3.84 (s, 6H); $^{13}$C NMR (126 MHz, $CDCl_3$): $\delta$ 161.1, 160.7, 102.7, 93.0, 89.4, 56.5.

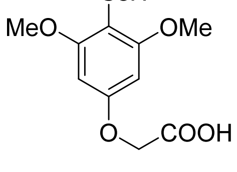


**14**

Intermediate **6** (150 mg, 0.58 mmol), methyl bromoacetate (133 mg, 0.87 mmol) and potassium carbonate (120 mg, 0.87 mmol) were dissolved in acetone (5 mL) and stirred at room temperature overnight. The reaction was quenched with water and the product was extracted with ethyl acetate. The organic layer was dried with sodium sulfate and vacuum concentrated. The resulted solid was dissolved in THF (5 mL) and NaOH (56 mg, 1.39 mmol) dissolved in $H_2O$ (3 mL) was added dropwise. After stirred at room temperature for 2 h, 0.5 mL 1 M HCl was added to quench the reaction. After vacuum concentrated to remove THF, the product in aqueous layer was purified via high-performance liquid chromatography (HPLC) to afford compound **14** (49 mg, 0.17 mmol, 29% yield) as a yellow solid. For HPLC purification (flow rate: 10 mL/min), solvent A is water containing 0.1% TFA while solvent B is acetonitrile containing 0.1% TFA. Solvent B percentage was increased gradient from 10% to 70% within 40 min. Compound **14** peak retention time on HPLC: ~ 34 min. $^1$H NMR (500 MHz, $CD_3CN$, mixture of monomer and dimer): $\delta$ 6.27 (s, 2H), 4.66 (s, 2H), 3.83 (s, 6H); $^{13}$C NMR (126 MHz, $CDCl_3$): $\delta$ 170.1, 162.7, 158.0, 102.7, 93.0, 92.3, 65.5, 56.8.

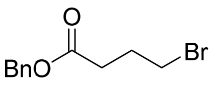


**7**

To a solution of 4-bromobutanoic acid (1.002 g, 6.0 mmol) in DCM (50 mL), oxalyl chloride (2 M in DCM, 3 mL, 6.0 mmol) and a drop of DMF (catalyst) were added. The mixture was stirred at room temperature for 2 h, then a solution of benzyl alcohol (683 µL, 6.6 mmol) and triethylamine (918 µL, 6.6 mmol) in DCM (5 mL) was added dropwise at 0 °C. After stirring at room temperature overnight, the reaction mixture was quenched with saturated aqueous solution of $NH_4Cl$, extracted with DCM, washed with brine, dried over anhydrous sodium sulfate and concentrated under vacuum. The crude mixture was then purified via flash column chromatography (hexane/ethyl acetate: 10/1) to afford compound **7** as a colorless oil (1.270 g, 5.0 mmol, 83% yield). $^1$H NMR (500 MHz, $CDCl_3$): $\delta$ 7.38-7.29 (m, 5H), 5.12 (s, 2H), 3.44 (t, $J$ = 6.5 Hz, 2H), 2.54 (t, $J$ = 7.5 Hz, 2H), 2.17 (qui, $J$ = 6.5 Hz, 2H); $^{13}$C NMR (126 MHz, $CDCl_3$): $\delta$ 172.4, 135.9, 128.6, 128.4, 128.3, 66.5, 32.8, 32.5, 27.8.

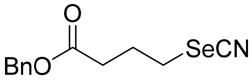


**8**

Benzyl 4-bromobutanoate (1.270 g, 5.0 mmol) and potassium selenocyanate (1.080 g, 7.5 mmol) were mixed in THF (20 mL) and stirred at room temperature for 24 h. Upon reaction completion, the mixture was diluted with water, and the product was extracted by ethyl acetate. The organic layer was washed with brine, dried over anhydrous sodium sulfate and concentrated under vacuum. The crude mixture was then purified via flash column chromatography (hexane/ethyl acetate: 5/1) to afford compound **8** as a yellow oil (1.273 g, 4.5 mmol, 90% yield). $^1$H NMR (500 MHz, $CDCl_3$): $\delta$ 7.39-7.31 (m, 5H), 5.13 (s, 2H), 3.09 (t, $J$ = 7.0 Hz, 2H), 2.56 (t, $J$ = 7.0 Hz, 2H), 2.23 (qui, $J$ = 7.0 Hz, 2H); $^{13}$C NMR (126 MHz, $CDCl_3$): $\delta$ 172.0, 135.6, 128.7, 128.5, 128.4, 101.2, 66.7, 33.0, 28.5, 26.0.

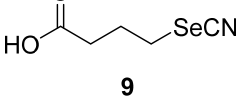


**9**

Lithium hydroxide (1 M aqueous solution, 5.4 mmol) was slowly added into a solution of compound **8** (1.273 g, 4.5 mmol) in THF (50 mL). The reaction mixture was stirred for 2 h and the byproduct was removed by extraction with ethyl acetate. Then the aqueous layer was acidified with 1 M HCl solution to pH ~3 and the product was extracted with ethyl acetate. The combined organic layer was dried over anhydrous sodium sulfate and concentrated under vacuum. The crude product **9** was used directly for the next step without further purification. Yellow oil (540 mg, 2.8 mmol, 62% yield). $^1$H NMR (500 MHz, $D_2O$): $\delta$ 3.09 (t, $J$ = 7.0 Hz, 2H), 2.50 (t, $J$ = 7.0 Hz, 2H), 2.23 (qui, $J$ = 7.0 Hz, 2H); $^{13}$C NMR (126 MHz, $D_2O$): $\delta$ 177.3, 105.7, 32.8, 28.7, 25.7.

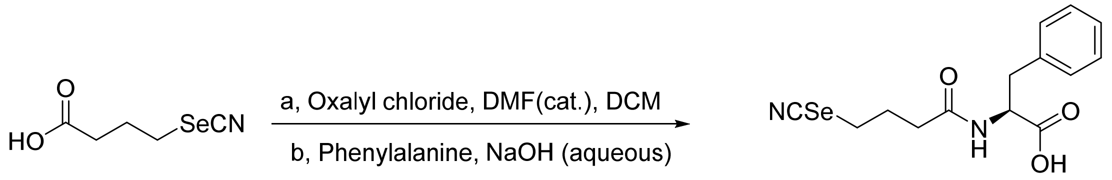


**9** **10**

To a solution of compound **9** (115 mg, 0.60 mmol) in DCM (5 mL), oxalyl chloride (2 M in DCM, 0.3 mL, 0.60 mmol) and a drop of DMF (catalyst) were added. The mixture was stirred at room temperature for 2 h and then add to the solution of phenylalanine (99 mg, 0.60 mmol) in NaOH aqueous (48 mg, 1.2 mmol) dropwise. After stirred at room temperature for 2 h, the reaction was quenched with 1 M HCl and the pH was adjusted to around 3. The product was extracted with ethyl acetate, dried over anhydrous sodium sulfate and concentrated under vacuum. The crude mixture was then purified via flash column chromatography (DCM/MeOH: 10/1, 1% acetic acid) to afford probe **10** as a peach solid (143 mg, 0.42 mmol, 70% yield). $^1$H NMR (500 MHz, $CD_3CN$ with 20% $CD_3OD$): $\delta$ 7.30-7.27 (m, 2H), 7.24-7.21 (m, 3H), 4.63 (dd, $J$ = 8.5, 5.0 Hz, 1H), 3.18 (dd, $J$ = 14.0, 5.0 Hz, 1H), 2.94-2.89 (m, 3H), 2.30-2.23 (m, 2H), 2.02 (quint, $J$ = 7.5 Hz, 2H); $^{13}$C NMR (126 MHz, $CD_3CN$ with 20% $CD_3OD$): $\delta$ 173.4, 172.8, 138.1, 130.1, 129.2, 127.6, 103.5, 54.2, 37.8, 35.2, 29.3, 27.7.

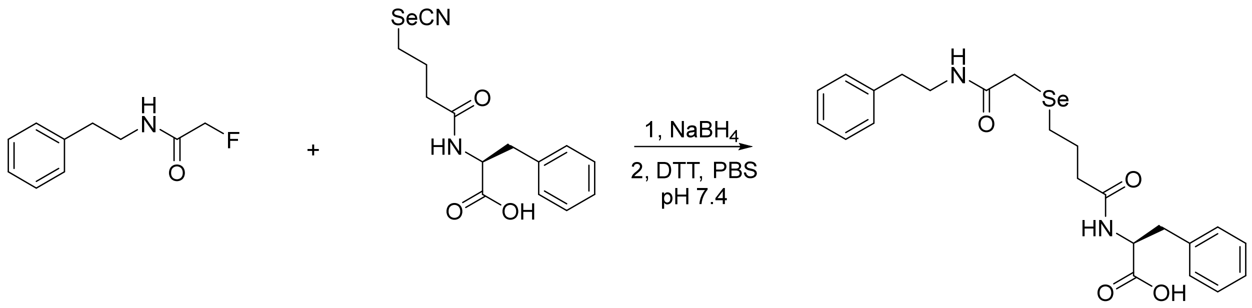


**11** **10** **16**

To a solution of compound **10** (84 mg, 0.25 mmol) in $CH_3CN$ (5 mL), $NaBH_4$ (57 mg, 1.5 mmol) aqueous solution (5 mL) was added. After stirred at room temperature for 10 min, DTT (370 mg, 2.4 mmol) and compound **11** (38 mg, 0.20 mmol) were added and the reaction mixture was stirred under nitrogen atmosphere overnight. Next day, the crude product was directly purified via high-performance liquid chromatography (HPLC). For HPLC purification (flow rate: 10 mL/min), solvent A is water containing 0.1% TFA while solvent B is acetonitrile containing 0.1% TFA. Solvent B percentage was increased gradient from 20% to 80% within 35 min. The extra selenol probe **13** peak retention time on HPLC: ~ 26 min; displacement reaction product **16** peak retention time on HPLC: ~ 28 min.

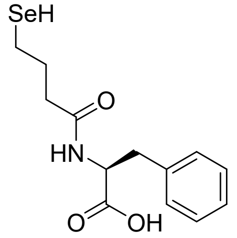

**13**

Compound **13** as a white solid (10 mg, 0.03 mmol). $^{1}$H NMR (500 MHz, $D_2O$ with 30% $CD_3CN$): $\delta$ 7.48-7.45 (m, 2H), 7.41-7.37 (m, 3H), 4.62-4.58 (m, 1H), 3.41-3.36 (m, 1H), 3.00-2.95 (m 1H), 2.70-2.61 (m, 2H), 2.42-2.31 (m, 2H), 2.20-2.17 (m, 2H); $^{13}$C NMR (126 MHz, $D_2O$ with 30% $CD_3CN$): $\delta$ 178.6, 174.7, 139.1, 130.1, 129.2, 127.3, 56.8, 38.7, 36.2, 28.3, 27.6.

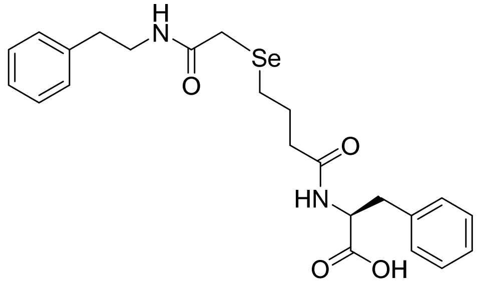

**16**

FSeDR product **16** as a white solid (62 mg, 0.13 mmol, 65%). $^{1}$H NMR (500 MHz, $D_2O$ with 30% $CD_3CN$): $\delta$ 7.26-7.21 (m, 4H), 7.19-7.14 (m, 6H), 4.56 (dd, $J$ = 9.5, 5.0 Hz, 1H), 3.34 (t, $J$ = 7.5 Hz, 2H), 3.12 (dd, $J$ = 14.0, 5.0 Hz, 1H), 3.00 (s, 2H), 2.87 (dd, $J$ = 14.0, 9.5 Hz, 1H), 2.72 (t, $J$ = 7.5 Hz, 2H), 2.31 (t, $J$ = 7.5 Hz, 2H), 2.12 (td, $J$ = 7.0, 1.5 Hz, 2H), 1.67 (qui, $J$ = 7.5 Hz, 2H); $^{13}$C NMR (126 MHz, $D_2O$ with 30% $CD_3CN$): $\delta$ 175.1, 175.0, 173.4, 140.0, 137.8, 130.1, 129.7, 129.4, 129.4, 127.8, 127.3, 54.5, 41.7, 37.6, 36.1, 35.5, 26.7, 25.4, 24.6.

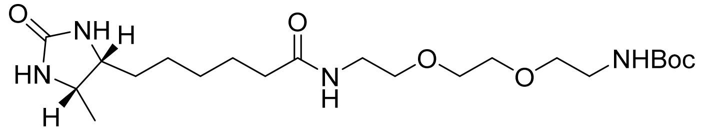

**18**

d-Desthiobiotin (214 mg, 1.0 mmol) was mixed with 1-[Bis(dimethylamino)methylene]-1H-1,2,3-triazolo[4,5-b] pyridinium 3-oxid hexafluorophosphate (HATU) (456 mg, 1.2 mmol) and DIPEA (209 µL, 1.2 mmol) in DMF (15 mL). After the mixture was stirred at room temperature for 10 min, t-Boc-N-amido-PEG$_2$-amine (298 mg, 1.2 mmol) was added. The resulting mixture was stirred at room temperature for 4 h, and then quenched by water. Ethyl acetate was added to extract the product from aqueous layer. The organic layer was dried with anhydrous sodium sulfate and concentrated under vacuum. The crude mixture was then purified via flash column chromatography (DCM/$CH_3OH$: 20/1) to afford compound **22** as a white solid (369 mg, 0.83 mmol, 83% yield). $^1$H NMR (500 MHz, $CDCl_3$): *δ* 3.76 (quint, *J* = 7.0 Hz, 1H), 3.66-3.62 (m, 1H), 3.56 (s, 4H), 3.53-3.48 (m, 4H), 3.40-3.37 (m, 2H), 3.30-3.20 (m, 2H), 2.15 (t, *J* = 7.5 Hz, 2H), 1.61 (quint, *J* = 7.5 Hz, 2H), 1.47-1.20 (m, 15H), 1.05 (d, *J* = 6.5 Hz, 3H); $^{13}$C NMR (126 MHz, $CDCl_3$): *δ* 173.3, 164.2, 156.1, 79.3, 70.22, 70.15, 70.06, 56.1, 51.4, 40.4, 39.2, 35.8, 29.5, 28.5, 25.8, 25.2, 15.8.

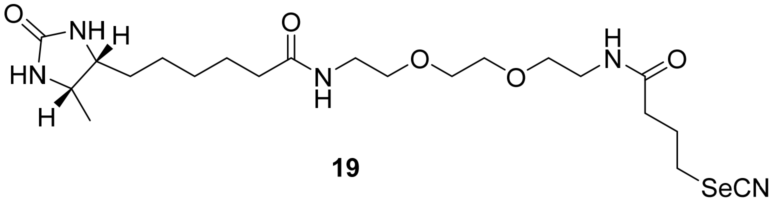


**19**

A solution of compound **22** (222 mg, 0.5 mmol) in DCM (5 ml) was treated with TFA (2 mL) for 2 h. The solvent was removed in vacuo and the crude was used directly for the next step without further purification. Compound **9** (97 mg, 0.5 mmol, 1 equiv) was mixed with HATU (228 mg, 0.6 mmol) and DIPEA (435 μL, 2.5 mmol) in DMF (5 mL). After stirred at room temperature for 10 min, the amine from last step (dissolved in 2 mL DMF) was added. After 4 h, the reaction mixture ws vacuum concentrated and the crude mixture was then purified via flash column chromatography (DCM/$CH_3OH$: 20/1) to afford compound **23** as a white solid (197 mg, 0.38 mmol, 76% yield). $^1$H NMR (500 MHz, $D_2O$): *δ* 3.82-3.76 (m, 1H), 3.69-3.64 (m, 1H), 3.56 (s, 4H), 3.51 (td, *J* = 5.5, 2.0 Hz, 4H), 3.28 (q, *J* = 5.0 Hz, 4H), 3.03 (t, *J* = 7.0 Hz, 2H), 2.34 (t, *J* = 7.0 Hz, 2H), 2.15 (t, *J* = 7.0 Hz, 2H), 2.08 (quint, *J* = 7.0 Hz, 2H), 1.50 (quint, *J* = 7.0 Hz, 2H), 1.43-1.37 (m, 2H), 1.29-1.15 (m, 4H), 1.00 (d, *J* = 6.5 Hz, 3H); $^{13}$C NMR (126 MHz, $D_2O$): *δ* 177.1, 175.2, 165.5, 105.6, 69.4, 68.86, 68.78, 56.0, 51.5, 38.9, 38.8, 35.6, 34.8, 28.6, 28.1, 26.8, 25.3, 25.2, 14.4.

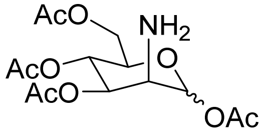


**20, $Ac_4ManNH_2$**

White solid (1.16 g, 67%). $^1$H NMR (500 MHz, $CDCl_3$): *δ* 6.20 (d, *J* = 8.0 Hz, 1H), 6.04 (d, *J* = 2.0 Hz, 1H), 4.99 (t, *J* = 10.0 Hz, 1H), 4.49-4.45 (m, 1H), 4.25 (dd, *J* = 12.0, 5.5 Hz, 1H), 4.20-4.16 (m, 1H), 4.04 (dd, *J* = 12.5, 2.5 Hz, 1H), 3.99-3.95 (m, 1H), 3.44 (d, *J* = 4.0 Hz, 1H), 2.14 (s, 3H), 2.12 (s, 3H), 2.08 (s, 3H), 2.07 (s, 3H); $^{13}$C NMR (126 MHz, $CDCl_3$): *δ* 172.3, 170.9, 170.8, 168.5, 92.0, 70.3, 69.1, 68.5, 62.6, 52.3, 23.3, 21.05, 21.00, 20.9.

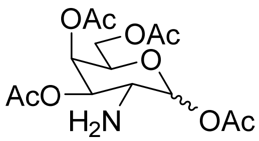


**21, $Ac_4GalNH_2$**

White solid (954 mg, 55%). $^1$H NMR (500 MHz, $CDCl_3$): $\delta$ 5.44 (d, $J$ = 8.5 Hz, 1H), 5.34 (d, $J$ = 3.0 Hz, 1H), 4.80 (dd, $J$ = 11.0, 3.0 Hz, 1H), 4.15-4.06 (m, 2H), 4.02 (t, $J$ = 7.0 Hz, 1H), 3.26 (t, $J$ = 9.5 Hz, 1H), 2.16 (s, 3H), 2.11 (s, 3H), 2.03 (s, 3H), 2.02 (s, 3H); $^{13}$C NMR (126 MHz, $CDCl_3$): $\delta$ 170.5, 170.4, 170.3, 169.3, 95.6, 77.2, 73.6, 71.7, 66.3, 61.3, 50.7, 21.1, 20.8, 20.8, 20.7.

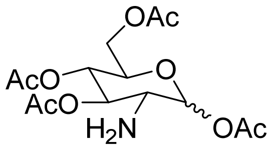


**22, $Ac_4GlcNH_2$**

White solid (1.30 g, 75%). $^1$H NMR (500 MHz, $CDCl_3$): $\delta$ 5.44 (d, $J$ = 8.5 Hz, 1H), 5.04 (t, $J$ = 9.5 Hz, 1H), 5.00 (t, $J$ = 9.5 Hz, 1H), 4.30 (dd, $J$ = 12.0, 4.5 Hz, 1H), 4.06 (dd, $J$ = 12.5, 2.0 Hz, 1H), 3.82-3.78 (m, 1H), 3.01 (t, $J$ = 9.5 Hz, 1H), 2.16 (s, 3H), 2.08 (s, 3H), 2.07 (s, 3H), 2.02 (s, 3H); $^{13}$C NMR (126 MHz, $CDCl_3$): $\delta$ 170.82, 170.81, 169.9, 169.4, 95.4, 75.2, 72.8, 68.3, 61.9, 55.1, 21.1, 20.9, 20.9, 20.8.

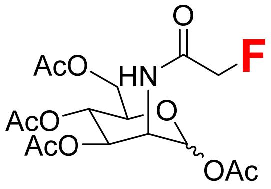


**23, $Ac_4$ManNFAc**

$^1$H NMR (500 MHz, $CDCl_3$, α:β = 2:1): $\delta$ 6.65 (dd, $J$ = 9.0, 3.0 Hz, 0.5H), 6.54 (dd, $J$ = 9.0, 3.0 Hz, 1H), 6.07 (d, $J$ = 2.0 Hz, 1H), 5.90 (d, $J$ = 1.5 Hz, 0.5H), 5.36 (dd, $J$ = 10.0, 4.0 Hz, 1H), 5.24 (t, $J$ = 10.0 Hz, 1H), 5.18 (t, $J$ = 10.0 Hz, 0.5H), 5.07 (dd, $J$ = 9.5, 3.5 Hz, 0.5H), 4.97-4.88 (m, 1.5H), 4.87-4.76 (m, 2H), 4.69-4.66 (m, 1H), 4.24-4.19 (m, 1.5H), 4.14 (dd, $J$ = 12.5, 2.5 Hz, 0.5H), 4.10 (dd, $J$ = 12.5, 2.5 Hz, 1H), 4.07-4.03 (m, 1H), 2.19 (s, 3H), 2.12 (s, 3H), 2.10 (s, 4.5H), 2.06 (s, 4.5H), 2.01 (s, 1.5H), 2.00 (s, 3H); $^{13}$C NMR (126 MHz, $CDCl_3$): $\delta$ 170.70 (α), 170.67 (β), 170.30 (β), 170.27 (α), 169.69 (β), 169.66 (α), 168.6 (β, $J$ = 17.8 Hz), 168.5 (β), 168.2 (α), 168.0 (α, $J$ = 17.8 Hz), 91.4 (α), 90.3 (β), 81.01 (β), 80.94 (α), 79.53 (β), 79.46 (α), 73.5, 71.6, 70.4, 69.0, 65.1 (α), 65.0 (β), 61.79 (α), 61.71 (β), 49.4 (β), 48.9(α), 21.0 (α), 20.91 (β), 20.87 (α), 20.82 (β), 20.80 (β), 20.77 (α), 20.74 (α); $^{19}$F NMR (471 MHz, $CDCl_3$): $\delta$ -224.64 (β anomer); -224.88 (α anomer).

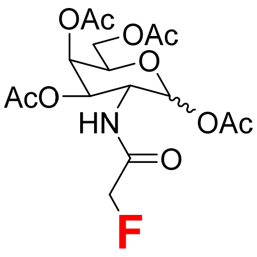

**24, Ac$_4$GalNFAc**

$^1$H NMR (500 MHz, CDCl$_3$, α anomer only): $\delta$ 6.40 (d, $J$ = 8.5 Hz, 1H), 5.81 (d, $J$ = 9.0 Hz, 1H), 5.40 (d, $J$ = 3.5 Hz, 1H), 5.22 (dd, $J$ = 11.0, 3.5 Hz, 1H), .4.71 (d, $J$ = 47.5 Hz, 2H), 4.42 (dd, $J$ = 20.0, 8.5 Hz, 1H), 4.20-4.10 (m, 2H), 4.07 (t, $J$ = 6.5 Hz, 1H), 2.17 (s, 3H), 2.13 (s, 3H), 2.05 (s, 3H), 2.02 (s, 3H); $^{13}$C NMR (126 MHz, CDCl$_3$): $\delta$ 170.60, 170.57, 170.3, 169.49, 168.2 ($J$ = 17.5 Hz), 92.7, 80.9 ($J$ = 186.9 Hz), 72.0, 70.2, 66.4, 61.4, 49.6, 21.0, 20.81, 20.80, 20.7; $^{19}$F NMR (471 MHz, CDCl$_3$): δ -224.65 (β anomer), -224.88 (α anomer).

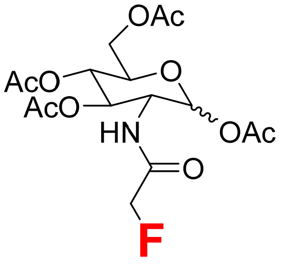

**25, Ac$_4$GlcNFAc**

$^1$H NMR (500 MHz, CDCl$_3$, α anomer only): $\delta$ 6.54 (dd, $J$ = 9.0, 3.0 Hz, 1H), 5.82 (d, $J$ = 9.0 Hz, 1H), 5.29 (dd, $J$ = 10.5, 9.0 Hz, 1H), 5.14 (t, $J$ = 9.5 Hz, 1H), 4.70 (dd, $J$ = 47.5, 3.5 Hz, 2H), 4.32-4.25 (m, 2H), 4.14 (dd, $J$ = 12.5, 2.0 Hz, 1H), 3.87-3.83 (m, 1H), 2.12 (s, 3H), 2.09 (s, 3H), 2.05 (s, 3H), 2.04 (s, 3H); $^{13}$C NMR (126 MHz, CDCl$_3$): $\delta$ 171.1, 170.8, 169.5, 168.18 ($J$ = 17.6 Hz), 92.2, 80.8 ($J$ = 186.9 Hz), 72.9, 72.3, 68.0, 61.7, 52.7, 38.7, 21.0, 20.9, 20.72, 20.70.

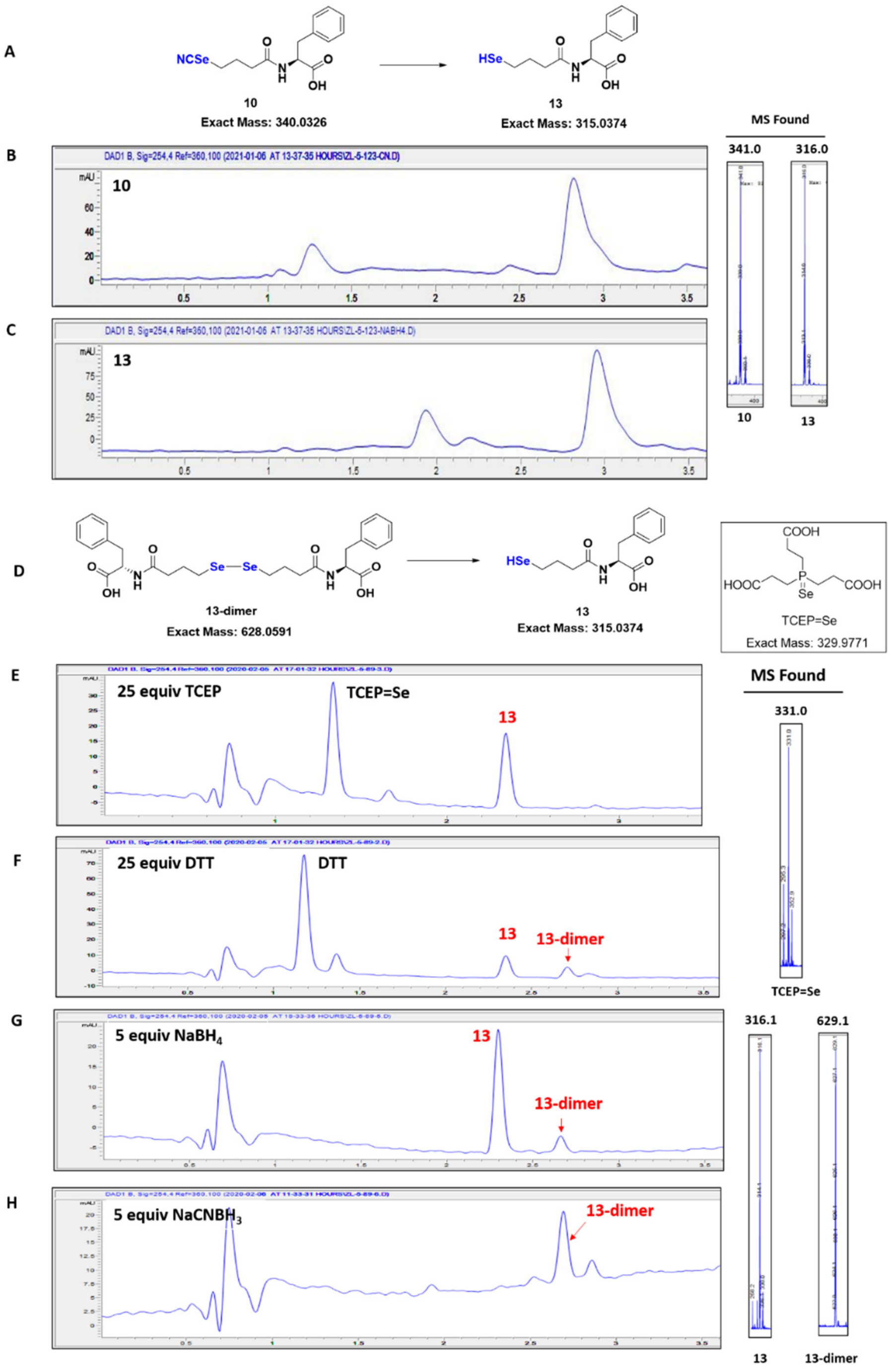


**Figure S1**. Deprotection of cyano-protected model selenol probe **10** and reduction of probe **13-dimer**. (A) Reaction scheme for cyano deprotection. (B) LC-MS spectra of cyano-protected model probe (**10**); MS (ESI) m/z calculated for $C_{14}H_{17}N_2O_3Se$ $[M + H]^+$: 341.0, found 341.0. (C) LC-MS spectra of deprotected model probe (**13**); MS (ESI) m/z calculated for $C_{13}H_{17}NO_3Se$ $[M + H]^+$: 316.0, found 316.0. (D) Reaction scheme for reduce diselenol probe **13-dimer**. (E-H) LC-MS spectra of **13-dimer** reduced by TCEP, DTT, $NaBH_4$, $NaCNBH_3$, respectively. **13-dimer** MS (ESI) m/z calculated for $C_{26}H_{33}N_2O_6Se_2$ $[M + H]^+$: 629.1, found 629.1; **13** MS (ESI) m/z calculated for $C_{13}H_{18}NO_3Se$ $[M + H]^+$: 316.0, found 316.1.
